# An Rtn4/Nogo-A-interacting micropeptide modulates synaptic plasticity with age

**DOI:** 10.1101/2022.01.26.477857

**Authors:** S. Kragness, Z. Clark, A. Mullin, J. Guidry, L.R. Earls

## Abstract

Micropeptides, encoded from small open reading frames of 300 nucleotides or less, are hidden throughout mammalian genomes, though few functional studies of micropeptides in the brain are published. Here, we describe a micropeptide known as the Plasticity –Associated Neural Transcript Short (Pants), located in the 22q11.2 region of the human genome, the microdeletion of which conveys a high risk for schizophrenia. Our data show that Pants is upregulated in early adulthood in the mossy fiber circuit of the hippocampus, where it exerts a powerful negative effect on long-term potentiation (LTP). Further, we find that Pants is secreted from neurons, where it associates with synapses but is rapidly degraded with stimulation. Pants dynamically interacts with Rtn4/Nogo-A, a well-studied regulator of adult plasticity. Pants interaction with Nogo-A augments its influence over postsynaptic AMPA receptor clustering, thus gating plasticity at adult synapses. This work shows that neural micropeptides can act as architectural modules that increase the functional diversity of the known proteome.

---

Schizophrenia (SZ) is a multifactorial disease that typically arises in late adolescence to early adulthood. The identification of molecular pathways that differentially influence brain function with adult maturation is key to understanding the age dependence of schizophrenia vulnerability. A microdeletion at chromosome 22q11.2 in humans, which causes the 22q11.2 deletion syndrome (22q11.2DS) conveys one of highest known genetic risks for schizophrenia, with ∼25% of 22q11.2DS patients developing SZ (1). Mouse models of the 22q11.2DS, carrying a deletion of the syntenic region on mouse chromosome 16, show robust, age-dependent phenotypes in multiple brain regions associated with SZ (2–6). For example, at and beyond 16 weeks of age, 22q11.2DS mouse models display increased long term potentiation (LTP) at hippocampal CA3-CA1 synapses that is associated with decreased performance in cognitive tasks (5). This serves as a sensitive cellular model for age-dependent onset of cognitive endophenotypes of SZ. While it is clear that multiple genes in the 22q11.2 region contribute to age-dependent hippocampal defects, not all of the genes involved have been identified.

Small open reading frames (sORFs) are newly-recognized genomic elements that encode for small peptides (SEPs/micropeptides) of ∼100 amino acids or less. sORFs are found throughout the genome (7), including in elements previously designated as non-coding RNAs (8–10), in antisense transcripts and in the assumed untranslated regions of larger mRNAs (11–14). Because of their simplicity, micropeptides make attractive candidates for elements that can be translated or degraded on the rapid timescale of brain activity. Further, because micropeptides lack complex structures that can affect multiple pathways, they may make for highly specific therapeutic targets. Molecular functions have been attributed to a few micropeptides in muscle (8), heart (15), and immune cells (16); however, few studies have examined micropeptides in the context of the adult brain. Recent work in Drosophila has revealed that the sight of a parasite can induce micropeptide expression in the adult brain, driving mating behavior (17). Further, translation of some sORFs has been shown to be upregulated in the brain with aging (18). We therefore asked whether micropeptides may contribute to age-dependent psychiatric disease associated with 22q11.2DS.

Recently, our laboratory described a novel sORF in the 22q11.2 disease-critical region that is conserved between mouse and human (19). The <u>P</u>lasticity <u>A</u>ssociated <u>N</u>eural <u>T</u>ranscript <u>S</u>hort (*Pants*) is upregulated at both the transcript and protein level in the hippocampus at 16 weeks of age, the stage of adult maturation at which neurophysiologic and behavioral phenotype changes were previously observed in models of 22q11.2DS (2–6). The age-dependent increase in Pants is particularly notable in the stratum lucidum, which houses the mossy fiber (MF) connections from the dentate gyrus to the region CA3 structurally-complex dendritic spines, known as thorny excrescences. Here, we show that the Pants peptide is localized both intracellularly at synapses and secreted into the extracellular space between synapses. We further show that Pants mutation exerts an age-dependent, overdominant, negative effect on synaptic plasticity at mossy fiber synapses. An endogenous proximity labeling screen revealed that Pants interacts with multiple intra- and extracellular proteins associated with memory and plasticity, including the Rtn4/Nogo-A protein. The Nogo-A pathway itself is an adult-specific negative regulator of plasticity through its effects on receptor clustering, and has previously been associated with SZ. Extracellular Pants at synapses quickly degrades with neuronal activity, resulting in increased AMPA receptor clustering. We therefore hypothesize that Pants is an adult-specific gate of Nogo-A plasticity regulation, and that activity-dependent degradation of Pants at select sites allows for plasticity at those synapses.

## 2. Results

### The *Pants^SOG-Y^* transgenic mouse line to study endogenous Pants

Previously, we showed in mice that Pants is expressed preferentially in areas CA3 and CA2 of the hippocampus, and that there is an upregulation of Pants peptide in these regions that is most prominent in the stratum lucidum between 8 and 16 weeks of age (19). However, as with other micropeptides studied (20), Pants is expressed at low levels and is poorly immunogenic, so antibody signal is low in most applications. In general, more traditional biochemical approaches to studying canonical proteins appear to be insufficient for tracking micropeptides. We therefore sought to verify our previous findings using a complementary, transgenic approach.

In order to obtain a more robust labeling of endogenous Pants, we used CRISPR/Cas9 to target the miniSOG and hemagglutinin (HA) tags to the C-terminus of the mouse *Pants* locus to generate a transgenic line that we refer to as *Pants^SOG-Y^* (**Fig. 1A**). Primers flanking the tags were used to genotype founders (**Fig.1B**), and specific detection of the Pants-miniSOG-HA fusion protein (predicted MW 26.32 KDa), was verified by immunoprecipitation with antibodies to HA, followed by western blotting for the HA tag (**Fig. 1C**). Using this approach, we verified that Pants levels increase in hippocampus between 8 and 16 weeks of age. We observed a 2.5-fold increase in Pants protein at 16 weeks compared to 8 weeks (n=3, P=0.004) (**Fig1D**), in agreement with previous findings.

**Fig. 1:**
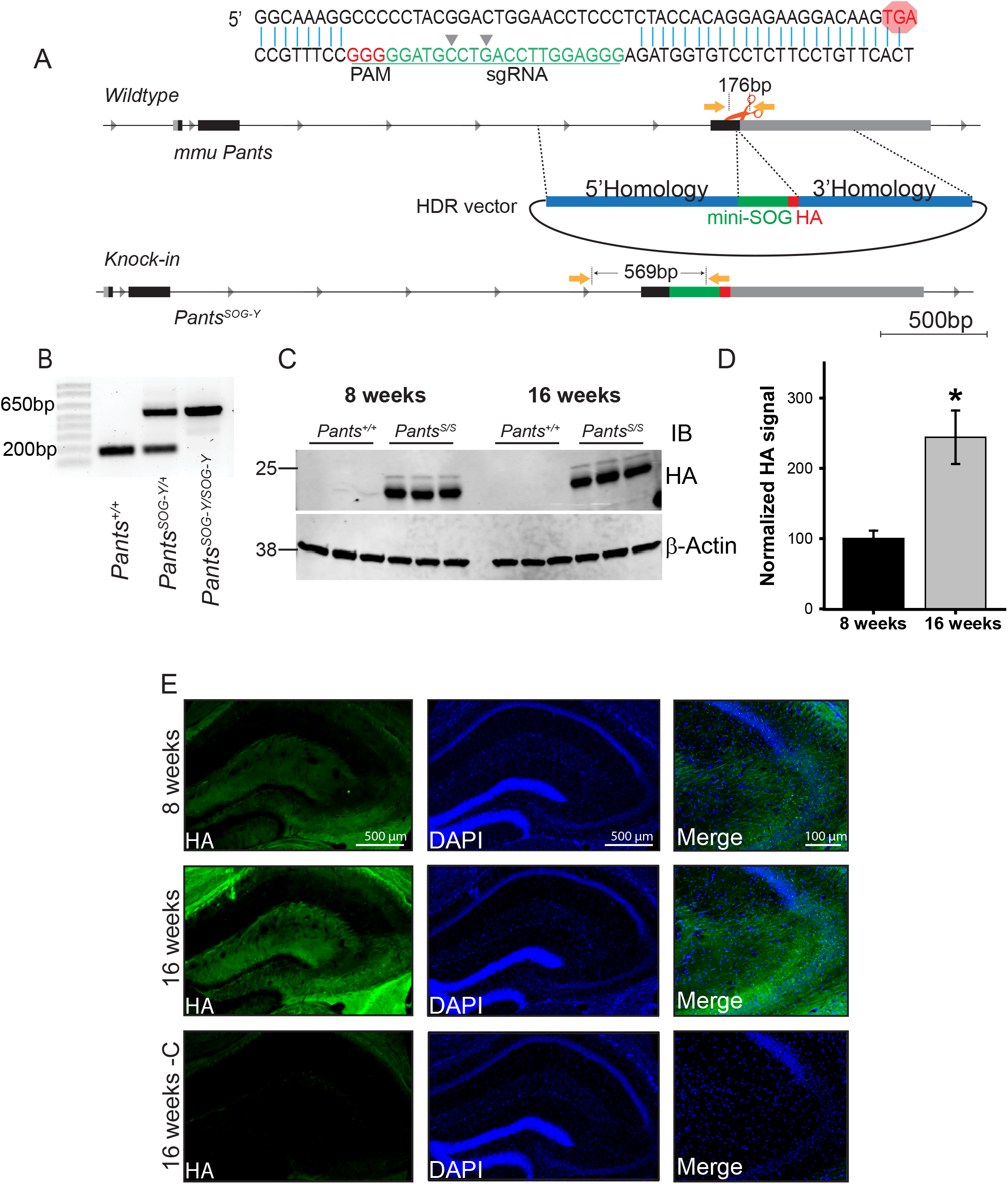
Generation and characterization of a transgenic mouse line to study endogenous Pants. **A)** CRISPR/Cas9 strategy to target the miniSOG and HA tags to the C-terminus of the endogenous *Pants* locus. **B)** Genotyping verification of insert in mouse tail biopsies. **C)** Quantitative Western blot of Pants immunoprecipitated from hippocampus of *Pants^SOG-Y^* and WT control mice at 8 and 16 weeks immunoblotted with HA antibody. **D)** Quantification of HA bands from (c), normalized to β-actin controls (n=3, P=0.004). **E)** Immunohistochemistry with HA antibody shows Pants localization in the hippocampus at 8 and 16 weeks.

Immunohistochemistry of 8- and 16-week-old *Pants^SOG-Y^* brain sections with the HA antibody was consistent with previous findings that the Pants peptide is most highly expressed in areas CA2 and CA3 in the hippocampus, with very little detectable CA1. Using this tagged approach, we see some staining in the molecular layer of the dentate gyrus, but not in the granule cell layer. We may be able to observe protein with the more sensitive HA antibody that was not detectable with the Pants antibody. Because staining in this region is excluded from cell bodies, the question remains as to the origin of Pants in this region of the DG. We have previously observed low-level staining of Pants in the cortex. Thus, this could represent perforant pathway expression. In areas CA3 and CA2, Pants is found around cell bodies of the pyramidal cell layer, and throughout regions containing mostly neural processes. In agreement with previous data, Pants levels increase at 16 weeks, with the most notable increase in the stratum lucidum (**Fig. 1E**).

### Pants localizes to synapses in hippocampal neurons

In order to study subcellular localization of Pants, we cultured primary neurons from *Pants^+/+^* control and *Pants^SOG-Y/+^* transgenic animals. Staining of DIV18-21 neurons with an anti-HA antibody followed by confocal microscopy showed that Pants is found in a punctate pattern throughout neuronal processes. Minimal background staining was observed in wildtype controls (**Fig. 2A**). Comparing Pants localization to that of the axonal marker Synapsin-1 and dendritic markers MAP2 and Shank-2 showed co-localization on both sides of the synapse (**Fig. 2A-C**). Overall, we observed approximately 70% of synapses to be positive for Pants (n=4, 100 synapses). Using both confocal microscopy and structured illumination microscopy (SIM), we often detected Pants staining that was located at or near the synapse, but which did not fully overlap with either pre- or post-synaptic markers (**Fig. 2B-C**, red arrows). This led us to ask whether Pants may be secreted from neurons.

**Fig. 2:**
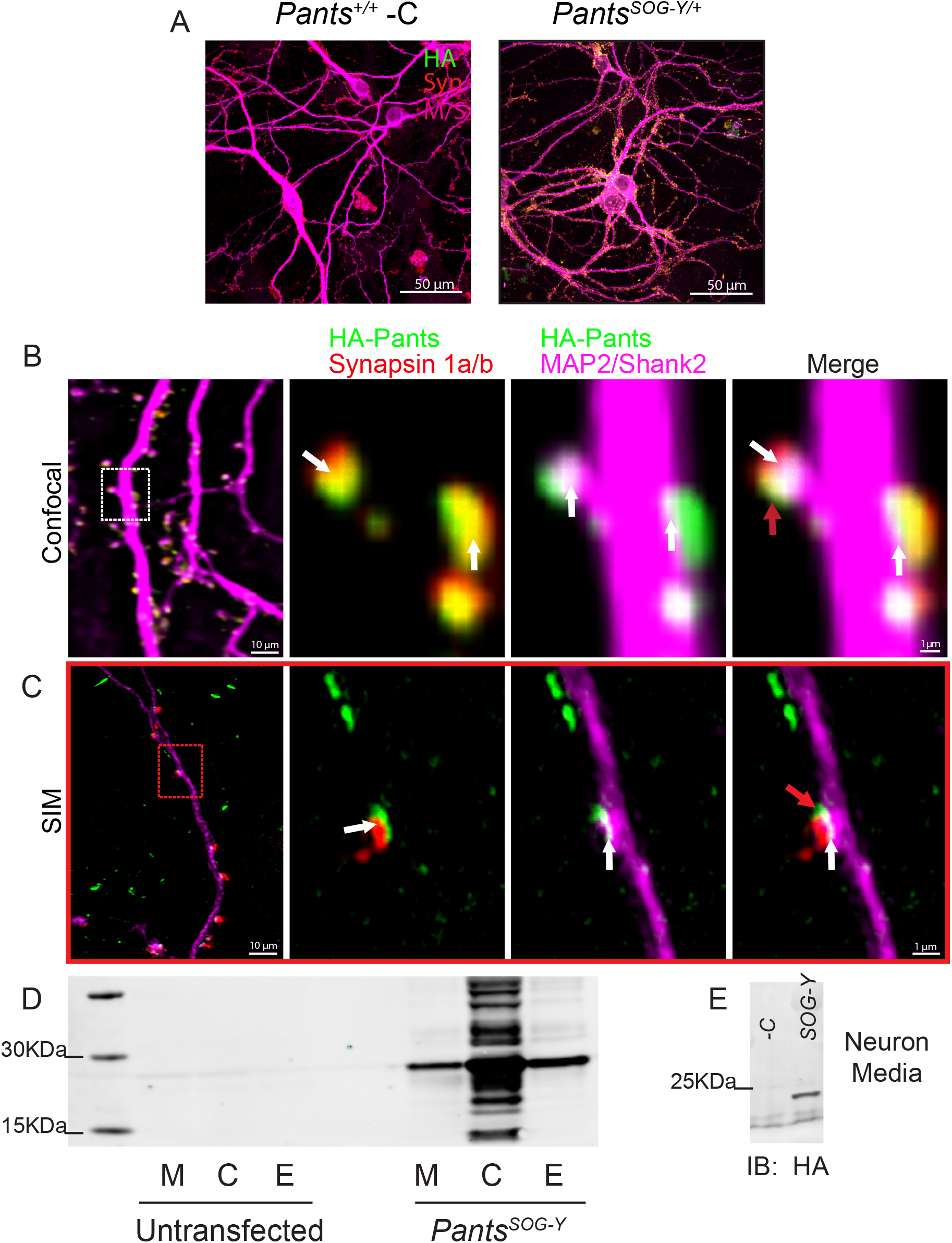
Synaptic localization and release of Pants. **A)** Immunocytochemistry with HA antibody shows Pants localization throughout cultured hippocampal neurons from *Pants^SOG-Y/+^* mice, but not wildtype controls. **B)** Immunocytochemistry of 18-21 div primary neurons from *Pants^SOG-Y^* hippocampus stained for HA, a Pre-synaptic marker (Synapsin 1a/b) and post-synaptic markers (MAP2 and Shank2) imaged with confocal microscopy. White arrows indicate areas of colocalization, and red arrows indicate suspected extra-synaptic pants that fails to co-localize with any markers. **C)** Structured Illumination Microscope imaging of the same staining from (A) to obtain better resolution. **D)** HA Western blot of HA IP from Nero2a cellular lysates (C), extracellular matrix (E) fractions, and proteins precipitated from cell media (M) after transfection with a construct for over-expressing the *Pants^SOG-Y^* fusion protein compared to untransfected controls. **E)** HA Western blot of HA IP of media collected from primary *Pants^SOG-Y^* hippocampal neurons or *Pants^+/+^* negative control neurons.

### Pants is secreted from Neuro2A cells and primary neurons

SecretomeP 2.0 (21) predicts the Pants peptide to be non-classically secreted with an NN-score of 0.808. In order to test whether Pants is secreted, we overexpressed a Pants^SOG-Y^ fusion protein in the mouse neuroblastoma cell line Neuro2A (N2a). We collected proteins released into fresh media over a 30-minute period. We then separated cellular proteins from extracellular matrix (ECM)-bound proteins. After immunoprecipitation in lysates from each fraction with an HA antibody to concentrate our fusion protein, we immunoblotted for the presence of HA. As shown in **Fig. 2D**, Pants is found in cells, and is detectable in the media and ECM fractions of cells overexpressing the Pants^SOG-Y^ fusion protein. To verify that these fractions had been correctly separated, we immunoblotted with antibodies specific for the ECM marker Aggrecan, and observed signal only in the ECM fraction (**fig. S2A**).

We further reasoned that Pants may be secreted from cells through large dense-core vesicles (LDCV), which is the route by which many neuropeptides are released. Using immunocytochemistry, we found modest co-localization with the LDCV-specific marker Chromagranin-B in *Pants^SOG-Y^* transgenic neurons (**fig. S2B**). While inconclusive, this raises the possibility that micropeptides may be secreted from neurons using routes similar to those used by more traditionally studied neuropeptides that are cleaved from larger proteins.

We next determined whether endogenous Pants might be released. To do this, we collected media from primary hippocampal neurons cultured from *wild-type* and *Pants^SOG-Y^* mice. As shown in **Fig. 2E**, HA-Pants was detected in the culture media collected from *Pants^SOG-Y^* neurons, but not from *wild-type* controls.

We further confirmed extracellular localization of Pants *in vivo*, in hippocampal sections from *Pants^SOG-Y/+^* brain. In adult (16-week) sections, Pants staining occurs in a similar punctate pattern as in primary neurons. A portion of these puncta co-localize with WFA, a marker for ECM glycoproteins (**fig. S4A**). Treating sections with chABC, an ECM-dissolving enzyme, Pants signal in area CA3 is reduced by 41.5% (**fig. S4B**), further confirming that Pants is partially extracellular and associates with ECM proteins in the adult brain.

Finally, we verified Pants secretion from neurons using an antibody feeding protocol that distinguishes extracellular from internal epitopes. This involves incubating live neurons with antibody to label extracellular epitopes prior to fixation, permeabilization and staining of intracellular Pants with an HA antibody raised in a different host (22). As shown in **Fig. 3A**, internal and extracellular Pants show little overlap, validating the approach. Further, extracellular Pants shows little overlap with pre-synaptic and post-synaptic markers. Meanwhile, intracellular Pants shows more overlap with pre-synaptic Synapsin than post-synaptic MAP2/Shank2 markers, as previously observed. **Fig. 3B** shows internal and extracellular Pants at morphologically distinct synapses. The first is a classical mushroom spine, and the second shows the morphological complexity of a maturing thorny excrescence. While CA3-CA3 connections occur on the former, the latter is the site in the intact hippocampus where DG-CA3 mossy fiber connections are formed. While we cannot draw analogies about the connectivity at these different spine types in culture, we observed both morphologically simple and complex synapses in our cultured neurons. Further, the complex synapses are positive for synaptoporin, a presynaptic marker for thorny excrescences (**Fig. S3**). We observed a similar pattern of internal and extracellular Pants staining at both types of spines. An illustration of localization at each type of synapse is shown in **Fig. 3C**. **Fig. 3D** shows the quantification of overlap using Mander’s colocalization coefficient in 190 spines for both internal and extracellular HA staining. As previously observed, intracellular Pants shows little overlap with extracellular staining, and overlapped more with presynaptic markers than postsynaptic markers. Meanwhile, the limited amount of extracellular Pants that does coincide with markers at all shows the most overlap with postsynaptic dendritic markers.

**Fig. 3:**
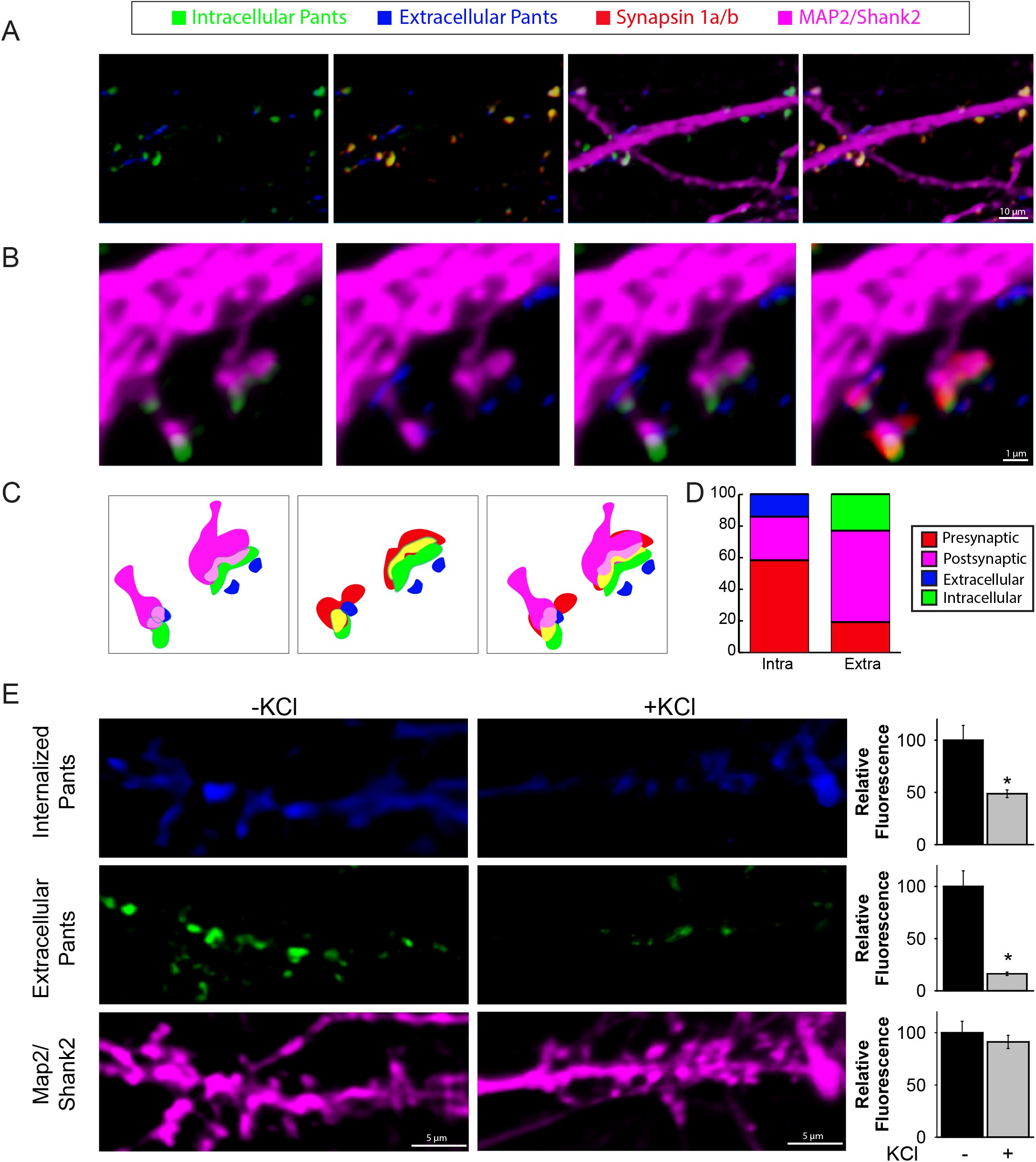
Extrasynaptic Pants is downregulated in response to neuronal activity. **A)** Contrasting extracellular/intracellular immunocytochemistry staining for HA in cultured hippocampal neurons. **B-C**) Co-localization of intra- and extra-cellular Pants with synaptic markers **D**) Quantification of Pants co-localization with pre- and post-synaptic markers correlation coefficients. **E)** Differentiated neuronal extracellular and internalized HA-Pants staining with and without 10 seconds 60mM KCl stimulation prior to fixation. Graphs at the right show quantification of HA fluorescence in dendritic ROIs, normalized to MAP2/Shank2 fluorescence. n=3 P<0.001(extracellular) and P=0.005 (internalized)P values generated by Mann-Whitney Rank Sum test.

### Extracellular Pants is rapidly degraded upon neuronal stimulation

We next asked whether Pants presence in the synapse was activity-dependent. To do this, we used a similar feeding protocol, but did not add an additional Pants antibody after fixation/permeabilization to stain intracellular Pants, but instead added a contrasting secondary antibody. This allowed us to distinguish extracellular Pants from Pants that was internalized during the course of the experiment. We performed this staining in the presence or absence of a brief (10 second) treatment with 60mM KCl, just prior to fixation. We observed a striking 83.7% reduction in extracellular Pants with KCl stimulation compared to unstimulated controls (n=3; P<0.001). We conclude that this reduction is likely due to proteolysis, rather than internalization, given that Pants internalization is decreased by 51.3% upon KCl treatment (n=3, P=0.005).

### Pants is an age-dependent, overdominant, negative regulator of mossy fiber LTP

Deletion of the region on mouse chromosome 16 that is similar to human chromosome 22q11.2 results in an age-dependent augmentation of LTP at Schaffer collateral synapses. Because Pants shows such striking age-dependent up-regulation in stratum lucidum and is located in this disease-critical region, we asked whether loss of Pants affects mossy fiber LTP in acute hippocampal slices at 8 weeks (prior to the onset of symptoms in 22q11.2 model mice) and at 16 weeks (after symptom onset). We confirmed that we were recording at MF synapses by performing frequency facilitation at 20Hz, observing a ratio of 2 or greater between the 3^rd^ and 1^st^ pulses (**Fig 4A**). Additionally, we observed a paired-pulse facilitation ratio of 1.75 or more for stimuli delivered at a 60ms interval (**Fig 4C**) (23, 24). At 8 weeks, in response to three 100Hz/1sec stimuli, we recorded indistinguishable LTP in area CA3 between *Pants^+/+^*, *Pants^+/-^*, and *Pants^-/-^* mice (n=6 mice/18 slices for *Pants^+/+^*, n=10 mice/30 slices for *Pants^+/-^*, n=5 mice/15 slices for *Pants^-/-^*; P>0.503). However, at 16 weeks, *Pants^+/-^* slices showed a robust (277%) increase in MF LTP, whereas *Pants^-/-^* LTP was indistinguishable from that of *Pants^+/+^* (**Fig. 4D**) (n=5 mice/18 slices for *Pants^+/+^*, n=5 mice/18 slices for *Pants^+/-^*, n=5 mice/19 slices for *Pants^-/-^*. P<0.001 *Pants^+/+^* vs. *Pants^+/-^*, P=0.157 for *Pants^+/+^* vs. *Pants^-/-^*, P<0.001 for *Pants^+/-^* vs. *Pants^-/-^*). We therefore conclude that Pants is an age-dependent, negative regulator of LTP, and that this effect is overdominant. After 1 hour of LTP recording, wash in of 1μM DCG-IV significantly reduced fEPSPs, a final validation that recordings were from the mossy fiber pathway (**Fig. S5**).

**Fig. 4:**
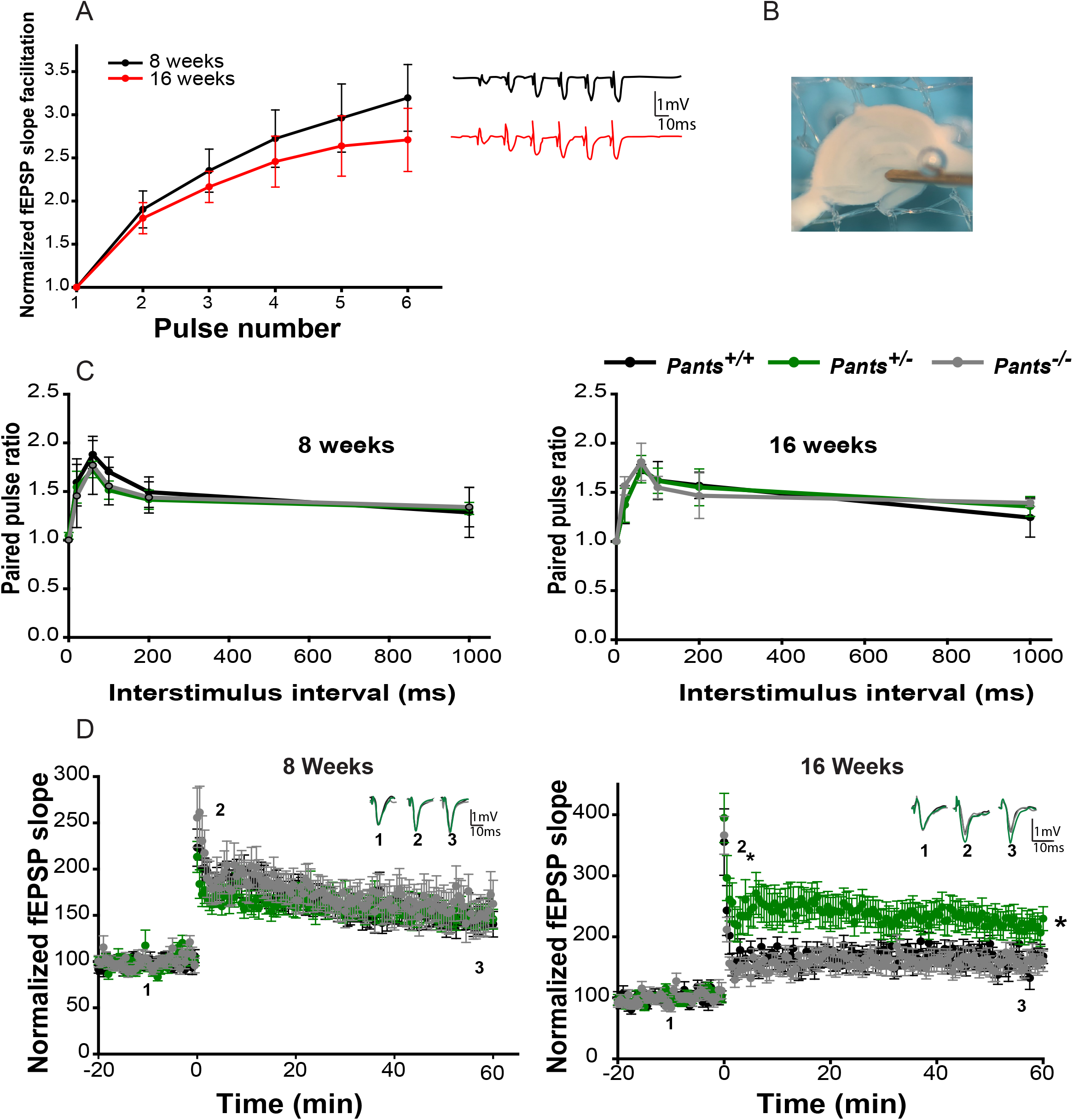
Pants exerts an age-dependent, overdominant, negative effect on hippocampal Mossy Fiber LTP. **A)** Frequency Facilitation at mossy fibers from acute hippocampal slices at 8 and 16 weeks of age (n=5). **B)** electrode placement for mossy fiber LTP measurement. **C**) Paired-pulse facilitation at MF synapses at 8 and 16 weeks of age (n=5). **D**) 100 Hz LTP recorded at MF synapses from 8 week old (left), and 16 week old (right) *Pants^+/+^*(black), *Pants^+/-^* (dark green), and *Pants^-/-^* (light green) acute hippocampal slices. 8 week (n=6 mice, 18 slices for *Pants^+/+^*, n=10 mice, 30 slices for *Pants^+/-^*, and n=5 mice, 15 slices for *Pants^-/-^*. All P > 0.503). 16 week (n=4 mice, 18 slices for *Pants^+/+^*, n=5 mice, 18 slices for *Pants^+/-^*, n=5 mice, 19 slices for *Pants^-/-^*. P<0.001 for *Pants^+/+^* vs. *Pants^+/-^.* P=0.157 for *Pants^+/+^* vs. *Pants^-/-^*. P<0.001 for *Pants^+/-^* vs. *Pants^-/-^*, Mann-Whitney).

### Pants interacts with proteins important for plasticity, cognitive disease, and aging

In order to ascertain the molecular mechanism by which Pants inhibits synaptic function, we sought to identify the proteins with which Pants interacts. Traditional affinity capture proteomics approaches are suboptimal when studying small, low-expression, and transient proteins, as they require large amounts of protein and stable interactions. Further, lysis artifacts can occur when spatially separated proteins are mixed during cell homogenization. Because Pants appears to be highly dynamic in neuronal cells, we used a proximity-labeling approach, fusing endogenously-expressed Pants with a biotin ligase, which transfers biotin to proteins that come within nanometers of the fusion protein. This approach has the advantages that labeling occurs in intact cells or tissues, eliminating lysis artifacts and allowing for the sensitive capture of transient and low expression targets using the high-affinity interaction between biotin and streptavidin.

The strategy for endogenous proximity labeling of Pants targets is outlined in **Fig. 5A**. We used the engineered biotin ligase TurboID, which is small (35 KDa) and has improved labeling kinetics over previous iterations of biotinylation enzymes (25). Using a similar strategy to that employed for the *Pants^SOG-Y^* transgenic mouse line, we generated transgenic N2a cell lines in which V5-tagged-TurboID was fused to the C-terminus of the endogenous *Pants* locus. The TurboID coding sequence was followed by the t2A self-cleaving peptide and GFP, so that properly targeted cells could be selected using green fluorescence. After co-transfection of the sgRNA/Cas9, and the HDR plasmid, we sorted GFP-positive single cells into the wells of 96-well plates. This allowed us to establish clonal cell lines to control for possible confounding results from Cas-9 off-targets. GFP-positive clones were expanded and genotyped for the correct targeting of TurboID to the *Pants* Locus (**Fig. 5B**). PCR products were then sequence verified. Western blotting of cell lysates using a V5-specific antibody showed a band of ∼48 KDa, the predicted size of the Pants-Turbo-ID fusion in the transgenic cell line, but not in control N2a cells (**Fig. 5C**).

**Fig. 5:**
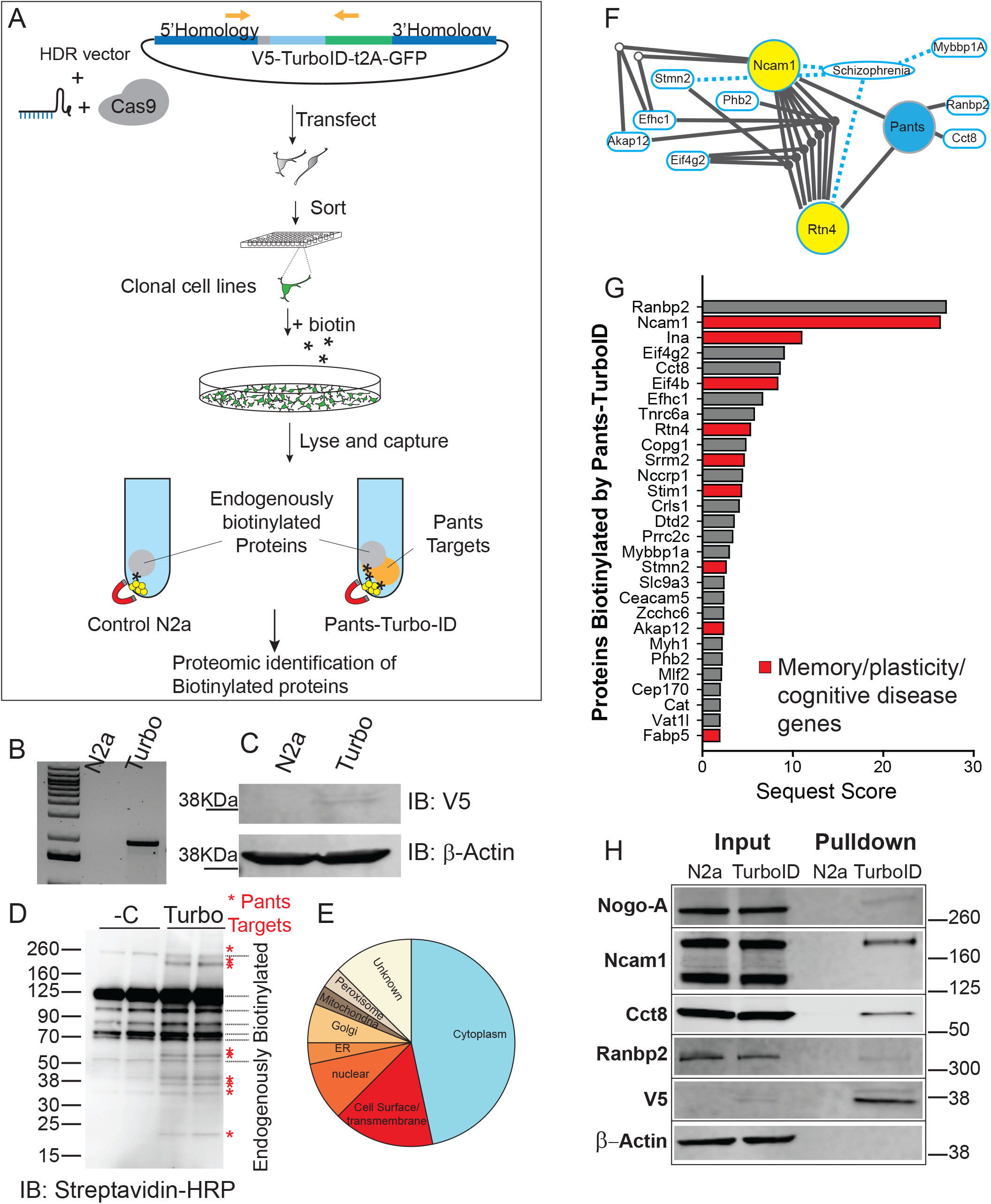
Endogenous proximity labeling proteomics reveals the Pants interactome. **A)** experimental design: a homomology-driven repair construct targeting the V5 epitope tag fused to the TurboID proximity labeling biotinylation enzyme followed by the self-cleaving t2A peptide and GFP to the 3’ end of the endogenous Pants locus was co-transfected with a vector carrying the sgRNA and Cas9 into Neuro2A cells. Single cells expressing GFP were then sorted into the wells of 96-well plates, generating clonal knock-in cell lines. **B)** Knock-in was verified by genotyping PCR of cell lines using a forward primer in the Pants gene and a reverse in GFP. **C)** Expression of the Pants-V5-TurboID fusion protein in cell lines was verified by Western blotting of cell lysates with an antibody to the V5 epitope. **D)** Treatment of Pants-Turbo cells with 50 μM biotin for 18 hours resulted in specific biotinylation of proteins not seen in N2A controls, as verified by Streptavidin-HRP immunoblotting. **E)** Pie chart showing localization pattern of Pants targets identified by proximity-labeling proteomics. **F)** IPA analysis of functional linkage between unique proteomic hits, showing SZ connections and inter-gene connections. **G)** Proteins uniquely biotinylated in Pants-TurboID cell line, ordered by Sequest Score. Those implicated in plasticity, memory or cognitive disease are highlighted in red. **H)** Validation of proteomics results by biotinylation followed by streptavidin pulldown and Western blot with selected antibodies specific to Pants targets identified by Mass spectrometry.

Cell lines were labeled by supplementing the cell culture media with 50μM biotin for 18 hours. Cells were collected and lysates were probed in western blots with a streptavidin antibody. As shown in **Fig. 5D**, N2a cells contain several endogenously biotinylated proteins, which likely correspond to mitochondrial carboxylases. In addition to these bands, samples from the Pants-Turbo transgenic cell lines contain additional, specific bands corresponding to Pants-interacting proteins. These specific bands were reproducible in multiple clonal cell lines, despite the high background of biotinylated proteins and low labeling due to the use of endogenous Pants. Next, we used streptavidin beads to capture biotinylated protein, and identified captured proteins by Mass Spectrometry (n=5 N2a control and 5 Pants-Turbo Tg samples). As expected, the most-represented proteins in the control cell line were carboxylases, which are known to be biotinylated (**table S3**).

**Table S2** and **table S4** show proteomic hits that were specifically captured in the Pants-TurboID cell line as compared to non-transgenic controls. As expected, the list is short, due to the above constraints. Localization data for proteomic hits from the UniProt database is summarized in **Fig. 5E**. The groups most represented are cytosolic and extracellular/transmembrane. This aligns with our imaging findings that Pants is both cytosolic and extracellular. Additionally, there were targets identified in ER and Golgi, which would be predicted for a secreted protein. IPA function analysis identified a number of proteins in the dataset associated with SZ, with the most functional connections between the proteins Ncam1 and Nogo-A. (**Fig. 5F**).

As highlighted in **table S2** and **Fig. 5G**, many of the Pants-binding proteins identified in this screen have been implicated in either plasticity or memory, including Ncam-1, α-internexin, Rtn4/Nogo-A, Stim1, Stathmin2 and Akap 12. In addition, a number of these proteins have been implicated in diseases of cognition.

To verify the hits from our proteomics screen, we performed biotin labeling experiments in a separate clonal Pants-Turbo transgenic cell line, and performed western blots on streptavidin-captured proteins, immunoblotting with commercial antibodies to selected protein hits from the screen. As shown in **Fig. 5H**, many of these proteins were verified to be specifically biotinylated in the Pants-TurboID line as compared to N2A controls. β-Actin was used as a negative control, and did not become biotinylated in the Pants-TurboID cells, whereas V5 labeling is prominent in this sample, indicating that Pants itself becomes biotinylated by TurboID. For each of the proteins tested, biotinylation occurred specifically in the Pants-Turbo sample, but not in N2a controls. Proteins verified included Rtn4/Nogo, Ncam-1, Ranbp2, and Cct8. In the case of NCam-1, we detected two of its isoforms, Ncam-140 and Ncam-180 in lysates; however, the Pants-TurboID transgene specifically led to biotinylation of the NCam-180 isoform (**Fig 5H**). This isoform is almost exclusively postsynaptic (26), and increases in spines following LTP induction in the perforant pathway (27), but it is not found at MF synapses (26). Thus, the interaction between Pants and Ncam-180 is likely not important for the plasticity effect of Pants at MF synapses. The 120KDa polysialated form of Ncam-1was not detected by our antibody, but it is important for hippocampal plasticity (28) and remains an important candidate.

### Pants interaction with Nogo-A at synapses enhances its inhibition of AMPA-receptor clustering

Rtn4/Nogo-A was an especially compelling hit from this proteomics screen for several reasons. First, like the 22q11.2 locus that includes *Pants*, the 2p16 genomic region that contains the *Rtn4* gene is a susceptibility locus for schizophrenia (29, 30). Additionally, the gene encoding one of its many receptors, *Rtn4R*, is localized to the 22q11.2 region. In organotypic slices, Nogo-A acts specifically on the structure and function of area CA3 of the hippocampus, where Pants is most highly expressed in adults. Nogo-A is well known for its role in axon guidance in the developing nervous system (31); however, in the adult nervous system, Nogo-A phenocopies Pants, acting as a negative regulator of synaptic plasticity. Particularly intriguing is the finding that the *Nogo-A* knockout has no effect on plasticity, whereas blockade with antibodies in acute murine slices (32) or knockdown with a miRNA approach in rats (33) results in an increase in LTP at Schaffer collateral synapses. This striking similarity with the Pants overdominant LTP increase lead us to question whether Pants may affect Nogo-A function at synapses.

To test this, we first asked whether Pants and Nogo-A interact at synapses. We cultured hippocampal neurons from *Pants^SOG-Y^* transgenic mice, and co-stained with antibodies to HA and Nogo. We observed significant overlap of Pants and Nogo at selected synaptic sites (**Fig. 6A**). This led us to wonder whether the interaction of Pants and Nogo may stabilize Nogo-A’s downstream signaling pathways. The synaptic impact of Nogo-A is well-studied, and is summarized in **Fig. 8**. The extensive extracellular domain of Nogo-A is divided into subdomains, each of which has been shown to interact with specific receptors. For example, the Nogo-66 domain interacts with a number of proteins, including Rtn4R (34). These interactions activate the RhoA pathway, stabilizing local actin dynamics in spines, thus blocking structural aspects of plasticity. The Δ20 domain is a disordered domain at the N-terminus of the Nogo-A protein that interacts with the Sphingolipid receptor S1Pr2, and it is this interaction that has been shown to dictate Nogo-A’s suppression of functional plasticity (35). Multiple laboratories have provided evidence that Nogo-A signaling through its receptors can affect receptor clustering at postsynaptic sites (36–38). We therefore hypothesized that Pants may structurally stabilize the Nogo-A interaction with receptors, to modulate postsynaptic AMPA-receptor clustering. To test this, we labeled membrane-localized GluR1 AMPA receptors by feeding a GluR1 antibody specific to an extracellular epitope, and measured AMPA fluorescence in spine membranes before and after KCl stimulation. With no KCl treatment, there is no clear relationship between Pants co-localization with Nogo-A and AMPA receptors (**fig. 5B-C**). However, after KCl stimulation, when extracellular Pants rapidly degrades, there is a strong negative correlation between co-localization of the remaining Pants with Nogo-A and AMPA fluorescence. That is, synapses where Pants remained were associated with decreased membrane AMPA fluorescence intensity compared to synapses lacking Pants associated with Nogo-A (**Fig. 6D-E**). We therefore conclude that Pants interaction with Nogo-A post-stimulus inhibits those synapses, whereas the rapid degradation of extracellular Pants at other sites creates a permissive environment for plasticity to occur.

**Fig. 6:**
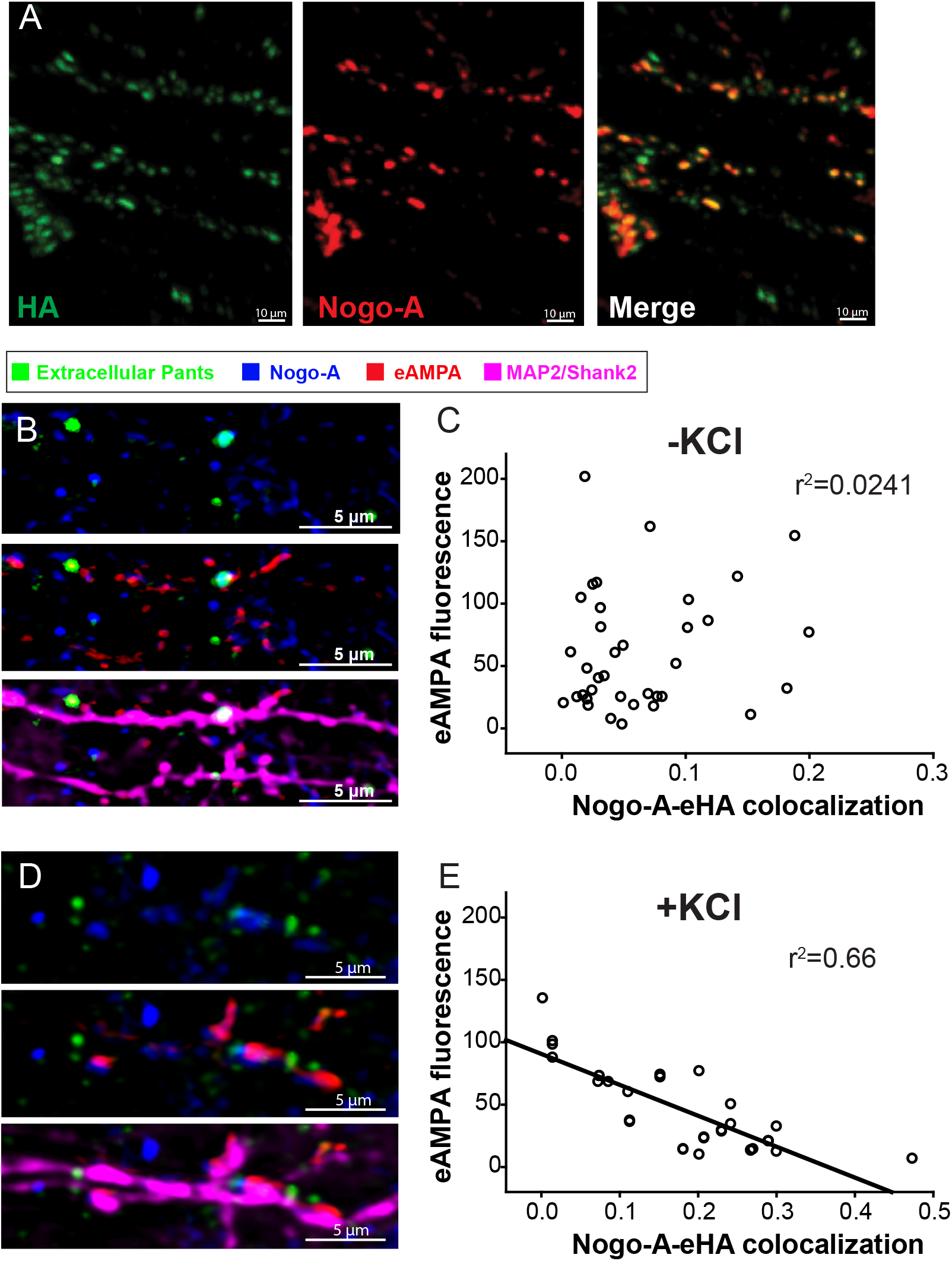
Pants interacts with Rtn4/Nogo in vitro and in vivo. **A)** Immunocytochemical co-localization of Pants and Nogo-A in cultured *Pants^SOG-Y^* hippocampal neurons **B)** AMPA receptor clustering at synapses where Pants colocalizes with Nogo-A versus Pants-free synapses in the absence of stimulation. **C)** Correlation between Pants-Nogo-A co-localization and GluR1 AMPA receptor fluorescence in the membrane of spine ROIs in the absence of stimulation. **D**) AMPA receptor clustering at synapses where Pants colocalizes with Nogo-A versus Pants-free synapses immediately after KCl stimulation. **E**) Correlation between Pants-Nogo-A co-localization and GluR1 AMPA receptor fluorescence in the membrane of spine ROIs immediately after KCl stimulation. n=3 samples, 40 ROIs each.

The previous results suggest that Pants augments Nogo-A-mediated inhibition of AMPA clustering. We therefore predicted that heterozygous Pants synapses might display increased AMPA receptor clustering. To study this, we cultured primary hippocampal neurons from *Pants^+/+^* and *Pants^+/-^* animals. We stained neurons for extracellular AMPA (eAMPA) and Nogo-A as above, along with a polyclonal antibody to Pants (19) and antibodies to Map2/Shank2 to label dendrites. We imaged synaptic ROIs associated with Nogo-A. As expected, we observed decreased Pants expression in *Pants^+/-^* neurons (**Fig. 7A-B**). Further, we verified the stimulation-dependent decrease in Pants in WT neurons (**Fig 7B**). In heterozyogous neurons, Pants associated with Nogo-A is significantly decreased compared to WT. However, the activity-dependent decrease in Pants is not significant in these neurons. We therefore conclude that, though Pants association with Nogo-A is reduced overall in heterozygotes, the interaction appears to be more stable with activity.

**Fig. 7:**
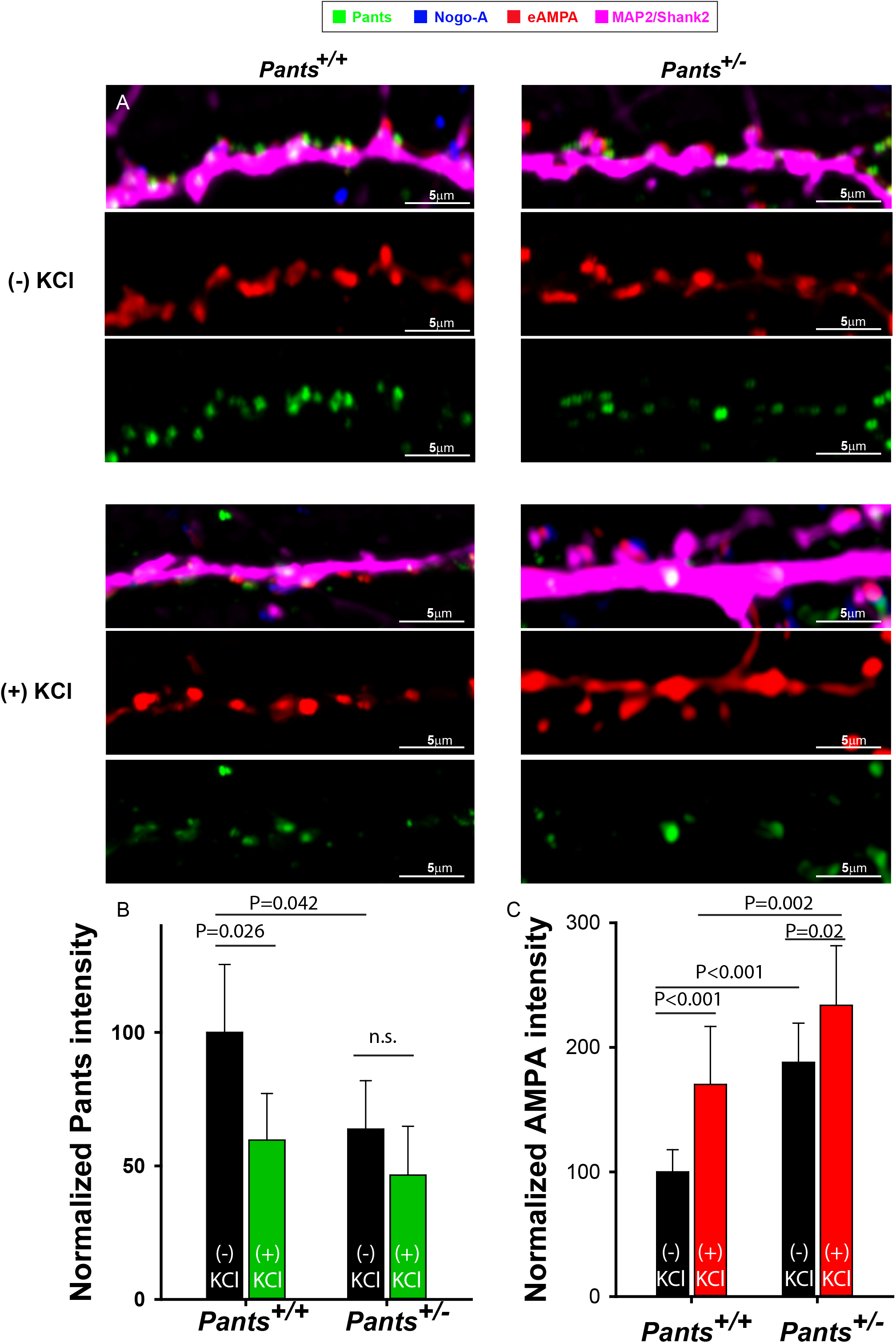
AMPA receptor clustering at spines increases in Pants +/- neurons. A) Representative dendrites from *Pants^+/+^* (left) or *Pants^+/-^* (right) hippocampal neurons stained with Nogo-A, an antibody to the extracellular domain of the GluR1 AMPA receptor, and a Nogo-A antibody. Staining was performed in the absence (top) or presence (bottom) of KCl stimulation. B) Quantification of Pants fluorescence intensity in neuronal processes C) Quantification of AMPA clustering at Nogo-A-associated spines in *Pants^+/+^* and *Pants^-/-^* neurons in the presence or absence of KCl stimulation. N=3 samples/ 30 ROIs, Mann-Whitney

**Fig. 8:**
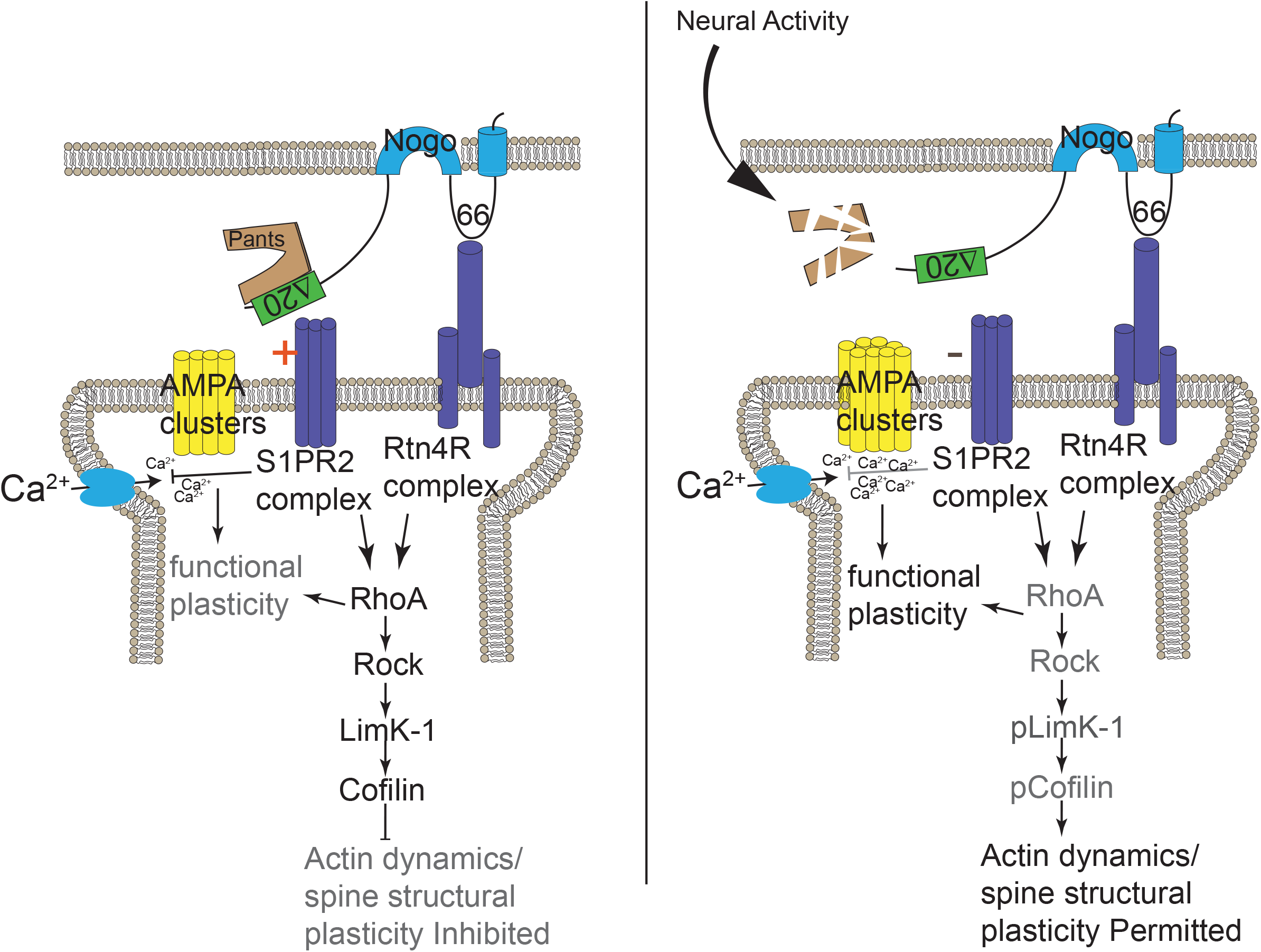
Model for the mechanism of Pants control of plasticity. Pants stabilizes the interaction of Nogo-A with receptors, inhibiting clustering of postsynaptic AMPA receptors at synaptic sites. Upon stimulation, Pants is degraded at synapses, reducing Nogo-A signaling and allowing AMPA receptor clustering at individual synapses.

We next tested whether AMPA clustering was increased in *Pants^+/-^* neurons. As shown in **Fig7A/C**, this is indeed the case. We observed an overall increase in the fluorescence intensity of AMPA clusters in *Pants^+/-^* neurons compared to WT. Further, while both genotypes showed an activity-dependent increase in AMPA clustering, this effect is blunted in *Pants^+/-^* neurons. This is consistent with the idea that, though there is less Pants associating with Nogo-A in the heterozyote, the interaction is more stable with activity. Thus, at *Pants^+/-^* synapses, we observe a baseline increase in AMPA clusters associated with Nogo-A, but a blunted activity-dependent increase in clustering. This indicates that reducing Pants primes some spines to be more responsive to glutamate signaling, in addition to allowing for increased AMPA clustering post-stimulus. A model for the Pants effects on the Nogo-A pathway is illustrated in **Fig. 8**. Understanding the difference in Pants-Nogo-A complex stability in the WT versus heterozygous conditions is therefore an important area for future investigation.

## 3. Discussion

While the juvenile brain is highly permissive to plasticity, the adult brain develops multiple mechanisms to gate plasticity, making the adult brain a more discerning network. For example, ECM remodeling from adolescence into adulthood plays a plasticity-dampening role in the adult brain (39). Expression patterns and levels of negative regulators of plasticity induction, such as Nogo-A pathway members, are altered with age, and these expression changes are associated with cognitive resilience in aged animals (40–42). Structural plasticity at spines has also been shown to change with age (43). Complex structural changes such as these are energetically costly for already taxed, aging neurons. Here we describe a mechanism by which the brain employs a micropeptide to rapidly modulate larger signaling complexes at the synapse. Micropeptides therefore represent a comparatively low-energy way for the brain to gate synaptic exchanges without the costly remodeling of larger protein complexes.

The 22q11.2 deletion syndrome results from a heterozygous microdeletion, emphasizing the importance of studying the heterozygous mutations of genes in this region. The overdominant effect of Pants on plasticity at MF synapses is particularly intriguing. Due to the low expression level of Pants, there is likely stoichiometric competition for Pants binding. Although Nogo-A is a negative regulator of plasticity, other proteins found bound to Pants, such as Ncam-1, have opposite effects on plasticity when blocked. Therefore, if Pants is partially reduced, competition between binding to negative and positive plasticity regulators may result in a net positive effect in plasticity. Our finding that Pants interactions with Nogo-A appear to be more stable in heterozygous neurons provide support for this idea. By contrast, complete loss of Pants eliminates competition, resulting in a balance of negative and positive effects of Pants target proteins. Further study of this phenomenon is ongoing.

The Pants interactome indicates functions for Pants in multiple subregions of neurons and synapses, but also potentially in other brain regions. For example, Vat1L is a marker for von Economo neurons (VENs) (44), which are most highly concentrated in socio-emotional cortical areas of human and closely-related primate brains. VENs show high expression of the DISC1 schizophrenia-associated gene (45), and the number of VENs is correlated with schizophrenia age of onset (46). This study confirms the importance of pursuing the human isoforms of Pants in both hippocampus and cortex.

The presence of micropeptides in the synaptic cleft represents a new frontier for neuropharmacology. While the major receptor systems targeted by more traditional therapeutics are pervasive throughout the brain, micropeptides may be employed in a more spatially and temporally restricted manner. Further study of these elements in normal health and disease therefore provides the potential for safer, more effective therapies.

## Materials and Methods

### Animals

Young (6-8 weeks) and mature (16-20 weeks) mice and their respective wild-type littermates of both sexes were used. Mice were generated and maintained on the C57BL/6 genetic background. *Pants^+/-^* mice were generated by KOMP (https://www.komp.org), using a velocigene targeting vector to disrupt the gene just downstream of the start codon. Mice were then crossed with EIIa-cre mice (Jackson Labs) expressing a germline specific Cre recombinase to delete the Neomycin cassette. Mice were genotyped using the following primers: PantsWTF: accatgtgaatctactgcctgaggg, PantsWTR: tatgtgggtgaatgcctgtagtccc, and PantsMF: gctcacctacactctgtgtatg and LacZR: gtgtagatgggcgcatcgtaac. PCR was performed using Go Green Taq (Promega) according to the vendor protocol. Cycling parameters were: 98oC 3min, followed by 35 cycles of 98°C 30 seconds, 53°C 30 seconds, 72°C 30 seconds, and a final 72°C extension for 5 minutes. This protocol produces a wildtype amplicon of 341 bp and a mutant amplicon of 718bp. *Pants^SOG-Y^* mice were generated using the Alt-R CRISPR-Cas9 crRNA kit (Integrated DNA Technologies), following the manufacturer’s protocol. Briefly, the crRNA sequence (5’ gggaggttccagtccgtaggggg 3’) was designed to target the area proximal to the stop codon in the Pants coding region using CHOPCHOP version 3 (47). This crRNA was duplexed with trans-activating crRNA (tracrRNA) and mixed with Alt-R S.p. HiFi Cas9 Nuclease (IDT) to form a ribonucloprotein complex. Complexes were electroporated into C57BL/6 zygotes with a double stranded donor DNA sequence. This dsDNA contained 1kb of homology upstream of the crRNA target sequence, the miniSOG and HA sequence added to the coding region of Pants just before the stop codon, followed by 1kb of downstream homology. Viable electroporated embryos at the two-cell stage were transferred to pseudopregnant females to create Pants^SOG-Y^ founders. Two positive males and two positive females were identified by genotyping and backcrossed with C57BL/6 mice to generate F1 mice. Primers used for genotyping were 5’ tgctcagaagcacaccttgg 3’ (forward) and 5’ aattagggctggaaatgctttgc 3’ (reverse) for the WT reaction and 5’ tgtgctaaaacagaacagttgcc 3’ (forward) and 5’ gaggttccagaatttcttcc 3’ (reverse) for the mutant reaction. PCR was performed using GoTaq Green (Promega) according to the vendor protocol. Cycling parameters were 98°C 3min, followed by 35 cycles of 98°C 30 seconds, 48°C for Mut reaction or 59°C for WT reaction 30 seconds, 72°C 30 seconds, followed by a final extension of 72°C for 5mins. Genotyping reactions result in a 176bp amplicon for the WT allele and a 569bp amplicon for the mutant allele. Amplicons were sequenced to verify the in-frame targeting of tags to the Pants locus. Potential off-targets predicted by the CHOPCHOP software were tested using PCR. No off-targeting was detected using this method (**Fig. S1**). Further, F1 animals were backcrossed 3 times to C57BL/6 mice to eliminate any unpredicted off-target mutations created by the CRISPR approach. The *Pants^SOG-Y^* strain can be obtained from Jackson Laboratory Repository (JAX Stock No. 037308). The care and use of animals were reviewed and approved by the Institutional Animal Care and Use Committee of Tulane University.

### Plasmid construction

To generate the donor DNA sequence used in the generation of the *Pants^SOG-Y^* mice, the upstream and downstream homology arms were amplified from mouse cDNA, adding the NheI and XbaI restriction sites for the upstream homology region using primers 5’ tatgctagctggcctggaactcagaaatcc 3’and 5’ tatctagacttgtccttctcctgtggtagagggaggttccaatcagtaggggg 3.’ XhoI and HindIII sites were introduced for the downstream homology region using primers, 5’ tatctcgagtaagcaactgcaaccctacc 3’ and 5’ ataaagcttaaaatacattaacaatgagg 3’. PCR products were digested, gel purified, and ligated into pcDNA3.1(-). The miniSOG-HA sequence was amplified from the miniSOG-C1 plasmid, introducing XbaI and XhoI sites using primers, 5’ tattctagaatggagaaaagttcgtgataac 3’ and 5’ atactcgagtcaagcgtaatctggaacatcgtatgggtaaccaccaccaccacctccatccagctgcactccg 3’, digested and ligated into pcDNA3.1(-), between the XbaI and XhoI sites of the homology arms. MiniSOG-C1 was a gift from Michael Davidson (Addgene plasmid #54821 ; http://n2t.net/addgene:54821 ; RRID:Addgene_54821). We modified the serine and arginine amino acids introduced by adding the XbaI site to an alanine and a glycine using the NEB Q5 Site-Directed Mutagenesis Kit (E0554S) altering the restriction site sequence from 5’ tctaga 3’ to 5’ gctgga 3’ (primers used were 5’ ccacaggagaaggacaaggctggaatggagaaaagtttcgtg 3’ and 5’ cacgaaacttttctccatggcggccgccttgtccttctcctgtgg 3’). To purify dsDNA fragment for electroporation, the newly generated plasmid was digested with NheI and HindIII to release the donor sequence and the fragment was purified using the Qiagen QIAEX II gel extraction kit.

To generate the plasmids used for the TurboID transgenic cell line generation, we introduced a NotI cloning site into the Pants homology plasmid described above, using site-directed mutagenesis and the following primers: 5’ ccacaggagaaggacaaggcggcggccgccatggagaaaagtttcgtg 3’ and 5’ cacgaaacttttctccatggcggccgccttgtccttctcctgtgg 3’. The miniSOG-HA sequence was digested out of the plasmid with NotI and XhoI, and V5-TurboID-t2A-EGFP sequence, amplified from the V5-TurboID-NES_pCDNA3 plasmid (Ting Lab, Addgene plasmid #107169 ; http://n2t.net/addgene:107169 ; RRID:Addgene_107169), was ligated into these sites. For *in vitro* expression of sgRNA and Cas9, we cloned the same sgRNA sequence from the generation of the Pants^SOG-Y^ mouse line into the pX330-U6-Chimeric_BB-CBh-hSpCas9, a gift from Feng Zhang (Addgene plasmid #42230 ; http://n2t.net/addgene:42230 ; RRID:Addgene_42230).

The Pants^SOG-Y^ overexpression (OE) plasmid was generated by amplifying the coding sequence for *Pants* from mouse cDNA using 5’ tatgctagcgccaccatggcggtcgcagggagc 3’ and 5’ tattctagatcacttgtccttctcctgtgg 3’ to introduce NheI and XbaI. The digested amplicon was fused to the miniSOG-HA sequence, amplified again from the miniSOG-C1 plasmid using the same primers used to clone miniSOG above. This full sequence was ligated into pcDNA3.1(-) to use for *in vitro* overexpression by transfection into N2a cells.

### Cell lines

N2A and transgenic cell lines were cultured in DMEM medium (SH30243, Cytiva) supplemented with 10% FBS (R&D Systems), GlutaMAX (Fisher Scientific), and Penicillin/Strepomycin (15140-122, Gibco). V5-TurboID-t2A-GFP was knocked into the endogenous Pants locus of Neuro2A cells by transfecting the TurboID donor sequence plasmid along with the Pants^SOG-Y^sgRNA plasmid. 48 hours after transfection, single EGFP positive cells were sorted into the wells of a 96 well plate using a Sony SH800 cell sorter to obtain clonal cell lines. Cell lines that were positive for GFP were expanded and genotyped using PCR. Primers used for genotyping were 5’ tgtgctaaaacagaacagttgccaa 3’ (forward) and 5’ aattagggctggaaatgctttgc 3’ (reverse) for the WT reaction and 5’ tgctcagaagcacaccttgg 3’ (forward) and 5’ ctcgagtctcaaccggtcttgtacagctcgtc 3’ (reverse) for the mutant reaction. PCR was performed using GoTaq Green (Promega) according to the vendor protocol. Cycling parameters were 98°C 3min, followed by 35 cycles of 98°C 30 seconds, 53°C 30 seconds, 72°C 30 seconds, followed by a final extension of 72°C for 5mins. PCR products were sequenced to verify correct targeting of the Pants locus. The expression of the transgene was verified by the detection of a 48 KDa band on a Western blot of cell lysates probed with an antibody to the V5 tag.

### Endogenous proximity labeling

Neuro2a cells (negative control) and Pants-TurboID N2a cells were grown to 90% confluency on 150 mm cell culture dishes. Biotin (100mM) in DMSO was added to a final concentration of 50μM to all samples for 18 hours. After labeling, plates were placed on ice and washed 5 times in 10ml ice cold PBS. Cells were displaced by scraping in 5ml PBS, collected by centrifugation at 5,000g at 4 °C, and frozen at -80 °C. Cell pellets were lysed in RIPA buffer (50mM Tris-HCl pH 7.5, 150mM NaCl, 1% Triton, 0.1% SDS, 0.5% Sodium Deoxycholate) with protease inhibitors (78428, ThermoFisherSci), and protein was quantified using a Protein 660 assay (22660, ThermoFisherSci). 2-4mg protein was added to 350μl streptavidin beads, and Streptavidin pulldown continued as previously described (48). For proteomics, beads were washed an additional 3 times with 1ml Tris-buffered saline to remove detergents, prior to on-bead trypsinization.

### On-Bead Trypsinization of biotinylated proteins

The procedure was adapted from (25). After several washes in 50mM Tris, pH 7.5 containing 2M Urea, beads and bound biotinylated proteins were incubated with 0.5ug trypsin in 50mM Tris, pH 7.5/2M Urea for 1 hour with shaking. This step was repeated for a second 15-minute period. These supernatants were pooled, the cysteines were reduced during a 1 hour incubation with 2ul of 500mM Tris(2-carboxyethyl)phosphine, and subsequently alkylated with a 30-minute incubation with 5ul of 375mM iodoacetamide in the dark. An additional 0.5ug of trypsin was added for an overnight incubation at room temperature.

The next day, samples were acidified by the addition of trifluoroacetic acid until 0.5%, and then trypsinized peptides were purified using C18 tips (Thermo). The eluted peptides were dried to completion until ready for LC-MS analysis.

### Liquid Chromatography-Mass Spectrometry (LC-MS)

The samples were run on a Dionex U3000 nano flow system coupled to a Thermo Fusion mass spectrometer. Each sample was subjected to a 90-minute chromatographic method employing a gradient from 2-50% acetonitrile in 0.1% formic Acid (ACN/FA) over the course of 65 minutes, from 50 to 99% ACN/FA for an additional 10 minutes, a hold step at 90% ACN/FA for 4 minutes and a re-equilibration into 2% ACN/FA. Chromatography was carried out in a “trap- and-load” format using EasySpray source (Thermo); trap column C18 PepMap 100, 5um, 100A and the separation column was EasySpray PepMap RSLC C18, 2um, 100A, 25cm. The entire run was 0.3ul/min flow rate. Electrospray was achieved at 1.9kV

MS1 scans were performed in the Orbitrap utilizing a resolution of 240,000, and data dependent MS2 scans were performed in the Orbitrap using high energy collision dissociation (HCD) of 30% using a resolution of 30,000 (49).

Data analysis was performed using Proteome Discoverer 2.4 using SEQUEST HT scoring. The background proteome was *Mus musculus* (SwissProt TaxID 10090, version 2017-10-15, downloaded on 01/19/2018). Static modification included carbamidomethyl on cysteines (=57.021), and dynamic modification of methionine oxidation (=15.9949), and biotinylation of N-termini and lysines (+226.0776). Parent ion tolerance was 10ppm, fragment mass tolerance was 0.02Da, and the maximum number of missed cleavages was set to 2. Only high scoring peptides were considered utilizing a false discovery rate (FDR) of 1%. Results were reported on grouped-replicate sample files and compared to identically treated control samples.

### Western blotting

Mice were sacrificed by decapitation, and hippocampi were dissected in cold PBS, frozen on dry ice and then stored at -80 °C. Lysates were prepared by needle passage in RIPA buffer containing protease inhibitors, followed by brief sonication. Lysates were clarified by centrifugation at 13,000g for 10 minutes at 4 °C. Protein concentrations were determined by a protein 660 Assay, diluted to equal concentration in sample buffer, and stored at -80 °C. Proteins were resolved on Novex 10-20% Tris-Glycine gels (Invitrogen, Carlsbad, CA) via SDS-PAGE and transferred to PVDF membranes (Invitrogen, Carlsbad, CA) according to manufacturer recommendations. Membranes were blocked with 3% BSA in TBST for 30 minutes at room temperature. Primary antibodies described in **table S1** were diluted in blocking buffer and incubated overnight at 4°C with gentle agitation. Membrane was washed 3x for 10 minutes in TBST. Secondary antibodies, goat anti-rabbit IgG IRDye 650 (1:10,000, LI-COR, Lincoln, NE) and donkey anti-mouse IRDye 800CW (1:15,000) were diluted in blocking buffer (LI-COR, Lincoln, NE) and incubated at room temperature in the dark on a shaker for 60 minutes. Membrane was washed 3x 10 minutes in TBST and then scanned on a LI-COR Odyssey scanner.

To verify specificity of labeling in Pants-Turbo cell lines, 40-50 micrograms of total cell protein was separated by SDS-PAGE gel electrophoresis, blocked overnight, and incubated with a steptavidin-horse radish peroxidase antibody (0.3micrograms/ml) for 1 hour prior to washing in TBST and ECL. To verify proteomic hits, labeled proteins were eluted from streptavidin beads by boiling for 10 minutes in SDS sample buffer containing 20mM DTT and 2mM biotin.

Samples were separated by SDS-PAGE and subjected to western blotting using antibodies specific to proteins identified in the proteomics screen. All western blots proceeded as described above, with the exception of Ranbp2, the antibody for which required blocking and antibody incubation in 5% non-fat dry milk in TBST.

### Immunohistochemistry

Fixed, cryoprotected, and OCT (VWR, Radnor, PA) embedded tissue was sectioned coronally at 25µm using a Leica CM3050S cryostat (Leica, Wetzlar, Germany). All tissue was then processed as follows. Tissue was permeabilized in 0.3% Triton X-100/1% BSA (Sigma, St Louis, MO)/1xPBS for 1 hour, washed in 1xPBS, and incubated with primary antibodies in 0.3% Triton X-100/1% BSA/PBS at 4°C overnight. Sections were washed 3x 5 minutes in 1xPBS and then incubated with secondary antibodies for 1 hour at room temperature. Secondary antibodies included donkey anti-rabbit AlexaFluor 488 (1:1,000; Life Technologies, Eugene, OR), goat anti-mouse AlexaFluor 568 (1:1,000; Life Technologies, Eugene, OR), and Hoeschst (ThermoFisher). Sections were washed again 3x 5 minutes in 1xPBS and mounted with Fluoromount-G (SouthernBiotech). Tissue or cells lacking the SOG-Y transgene were used as a negative control. Images were obtained with a 10x or 20x objective on a Nikon A1 confocal microscope.

### Primary culture of hippocampal neurons

Neuronal cultures were obtained from postnatal day 0 (P0) to P2 C57BL/6 mouse hippocampi from both sexes. Hippocampi were removed and digested with papain (2mg/mL, Worthington) in Hibernate without Calcium (BrainBits) and 0.1% DNase at 37°C for 10 minutes. Following titration of tissue by sterile Pasteur pipet, dissociated cells were re-suspended in culture media (Neurobasal A media (ThermoFisherSci) supplemented with GlutaMAX, pen-strep, B27, 1mM HEPES, and 10% horse serum) and plated at 30,000 cells/cm^2^ on poly-d-lysine and laminin coated dishes. Cells were maintained in feeding media (Neurobasal A media (ThermoFisher) supplemented with GlutaMAX, pen-strep, B27, and 1mM HEPES) by half media changes every 3-4 days until DIV14-21. Cultures were incubated at 37°C, 5% CO^2^ and 95% relative humidity.

### Immunocytochemistry

Cultured cells were fixed in 4% PFA for 10mins and washed 3x with PBS. Followed by blocking in 0.3% Triton X-100/5% BSA in PBS for 1 hour. Cells were then incubated in primary antibody overnight at 4°C with agitation. The next day, cells were washed 3x with PBS and incubated with secondary antibodies for 1 hour at room temperature. Cells were washed again 3x with PBS before allowing to dry and mounting with Fluoromount-G (SouthernBioTech).

Differential labeling of intracellular and extracellular HA epitopes protocol was adapted from (22). Briefly, the mouse anti-HA antibody (Santa Cruz) was added to primary neuron media and incubated for 2 hours at 37°C to allow for primary antibody binding to extracellular epitopes. Neurons were then fixed in 4% PFA for 5mins at room temperature followed by 3X washes with PBS. Cells were then blocked for 30mins with 5% BSA in PBS at room temperature. A fluorescently labeled secondary antibody (donkey anti-mouse 488, Invitrogen) was then applied to neurons for 1 hour to specifically label the extracellular HA epitopes. Any protein that was internalized after primary antibody binding would not be labeled with this secondary. Cells were then washed 3x with PBS. Further blocking was conducted on unpermeabilized neurons by incubation with an unlabeled AffiniPure F_ab_ fragment Donkey anti-Mouse IgG (H+L) (Jackson ImmunoResearch) at 0.13mg/mL overnight at 4°C. The next day, neurons were washed 3x 5min with PBS followed by a post-fix in 4% PFA in PBS for 5min at room temperature. After 2X 5min washes with PBS to remove the fixative, neurons were permeabilized and blocked in 5% BSA in PBS with 0.1% Triton-X-100 for 30mins at room temperature. Permeabilized neurons were then incubated in primary antibody solutions overnight at 4°C with agitation. The mouse anti-HA antibody was not introduced to the neurons again during this incubation. The next day, neurons were washed 3x 5mins in PBS before incubation with secondary antibodies for 1 hour at room temperature. This incubation included a 405-conjugated donkey anti-mouse secondary antibody to differentially label the intracellular HA epitopes. Neurons were washed for a final 3x 5mins in PBS before mounting with Fluoromount-G (SouthernBioTech) and allowed to cure overnight at 4◦C before imaging with a Nikon C1 confocal microscope with 60X objective.

### Electrophysiology

400 micron acute, transverse hippocampal slices were prepared in cold dissection buffer (125mM choline chloride, 2.5mM KCl, 0.4mM CaCl_2_, 6mM MgCl_2_, 1.25mMNaH_2_PO_4_, 26mM NaHCO_3_, and 20mM glucose (285-295mOsm)) as previously described (6). Slices recovered for 1hour in ACSF (125mM NaCl, 2.5mM KCl, 2mM CaCl_2_, 2mM MgCl_2_, 1.25mM NaH_2_PO_4_, 26mM NaHCO_3_, and 10mM glucose) and were continuously superfused with warm ACSF (30-32 °C) at a rate of 2-3ml/minute for recording. All solutions were equilibrated by continuous bubbling with 95% O_2_/5%CO_2_. Field EPSPs were recorded from the stratum lucidum in area CA3 using an extracellular glass pipette with a resistance of 3-5M Ω filled with ACSF. Mossy fibers were stimulated with a bipolar tungsten electrode placed in the granule cell layer of the dentate gyrus (**Fig. 4B**). MF stimulation was confirmed by 3 criteria: 1) the ratio of paired-pulse facilitation at the 60ms interval was ≥1.75; 2) frequency facilitation at 20Hz had a ratio of ≥2.0 as determined by the third response compared to the first response; and 3) bath application of 1µM DCG-IV (0975, Tocris) at the end of LTP recording significantly reduced the evoked fEPSPs. Stimulation intensities were chosen to produce a fEPSP of 30% of maximium reponse. For LTP recordings, baseline was monitored for 20 minutes, prior to LTP induction by three, 1-second trains of 100Hz, delivered at an interval of 10 seconds (50).

### Enzymatic degradation of ECM

100 micron acute, transverse hippocampal slices were obtained as above. Slices were incubated in recording aCSF with 0.2U/mL chABC (C3667, Sigma-Aldrich) or control recording aCSF for 1 hour at 37 ◦C with agitation. After 1 hour, slices were fixed in 4% PFA for 10min and washed 3x in PBS. Slices were mounted on slides and processed using IHC methods described above. WFA lectin (L-1516, Sigma-Aldrich) staining was used to monitor the extent of ECM degradation.

### *In vitro* ECM analysis

Detection of Pants^SOG-Y^ protein in media, cell bodies, and ECM was adapted from a previously established protocol (51). N2a cells were transfected with the Pants^SOG-Y^ overexpression plasmid and allowed to grow for 48 hours. Media on cells was completely exchanged for serum free media, 30 mins before harvesting. Serum free media from overexpression and untransfected control cells was collected and proteins were later precipitated using trichloroacetic acid (TCA). In brief, 0.11 volumes of ice cold 100% TCA was added to protein samples and incubated on ice for 20 minutes. After incubation, samples were spun at 14.8xg for 10 minutes at 4°C. Supernatants were removed and pellets were washed with ice cold 100% acetone and re-centrifuged. Supernatant was again removed and pellets were allowed to dry on ice for 10 minutes. Protein pellets were reconstituted in RIPA buffer. To specifically detach cell bodies from ECM, 20mM ammonium hydroxide was added to culture dishes and allowed to incubate for 5 mins at room temperature with gentle agitation once per minute. Cell body material was later TCA-precipitated by using the same method described for media fractions. To harvest ECM, culture dishes were washed 3x with de-ionized H_2_0, and the absence of remaining cells was confirmed under a microscope prior to mechanically lifting ECM material from the dish in RIPA using a cell scraper. All three fractions for overexpression and untransfected cells were immunoprecipitated using anti-HA magnetic beads (ThermoFisherSci) and the presence of HA-Pants was confirmed by Western blot.

### Imaging and Statistical Analysis

PClamp (Version10.6, Axon Instruments) was used for acquisition and analysis of fEPSPs. All confocal images were run through denoise and deconvolution processing in Nikon Elements software version 5.30.03. Elements was also used for colocalization analysis on selected ROIs in intra/extracellular HA labeling experiments in neurons. All statistical analysis for confocal images and electrophysiology data was done in SigmaPlot (Version 13.0, Systat Software Inc.). To determine negative correlation between Nogo-A/Pants colocalization and AMPA intensity, we used the Mander’s colocalization coefficients to determine the degree of co-localization between Pants and Nogo-A at synapses after stimulation. This data was then plotted against the intensity of AMPA signal at the respective spines and a linear line was fit to the plot. Data are all represented as means+/- standard error. Both Male and Female animals were used. All data was analyzed by Mann-Whitney Rank Sum test with significance set at p<0.05.

## Acknowledgments

The authors would like to thank Dr. Charles Nichols, Jarmell McGill, Dr. Amy Pierce, and Provost Dr Robin Forman for helping to save this research after hurricane Ida.

## Funding

A NARSAD Young Investigator Award from the Brain and Behavior Research Foundation. (LRE)

Louisiana Board of Regents RCS LEQSF(2018-21)RD-A-16 (LRE)

Tulane University Carol Lavin Bernick Faculty Grant (LRE)

Tulane Brain Institute Marko Spark Fund Award (LRE)

## Author contributions

Conceptualization: SK, LRE

Methodology: SK, ZC, AM, JG, LRE

Investigation: SK, ZC, AM, JG, LRE

Visualization: SK, LRE

Funding acquisition: LRE

Project administration: LRE

Supervision: SK, LRE

Writing – original draft: SK, LRE

Writing – review & editing: SK, ZC, AM, JG, LRE

## Competing interests

The authors declare that they have no competing interests.

## Data and materials availability

All data are available in the main text or the supplementary materials.

## Notes

### Competing Interest Statement

The authors have declared no competing interest.

### Summary of Updates

AMPA clustering in Pants heterozygous animals was measured. Controls to verify mossy fiber recordings were added

