## Supplementary for "An Rtn4/Nogo-A-interacting micropeptide modulates synaptic plasticity with age": PLOS_Supplement_Kragness2022.docx

**This PDF file includes:**

Figs. S1 to S5

Tables S1 to S2

Captions for Data tables S3 to S4

**Other Supplementary Materials for this manuscript include the following:**

Supp_Table_S3

Supp_Table_S4

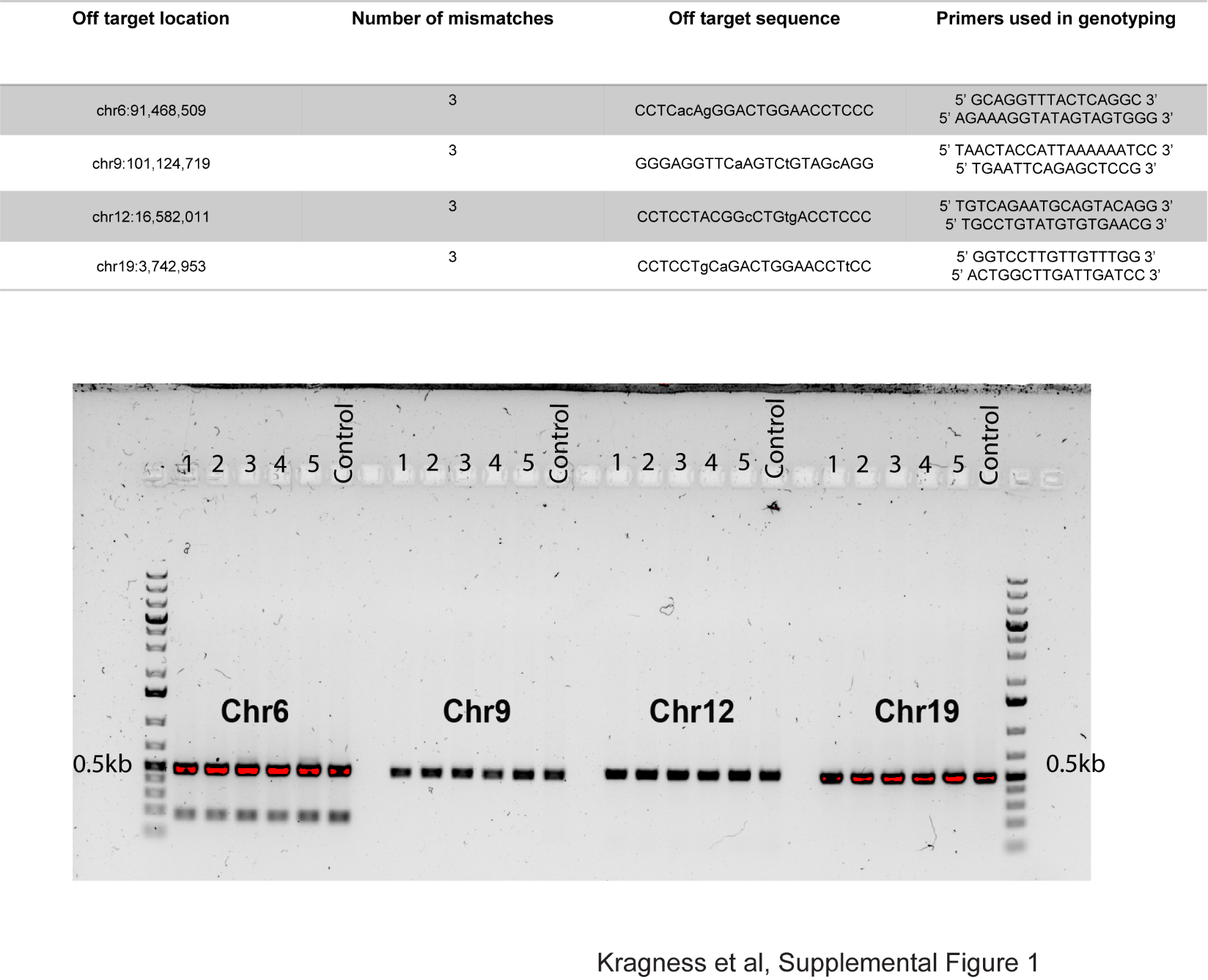

**Fig. S1: Test of *Pants^SOG-Y^* founders for CRISPR/Cas9 off targeting**. Above, Predicted potential off-target genes for our CRISPR HDR strategy. Primers were designed up and downstream of the target site to produce a 0.5Kb band for detection of the *Wildtype* locus. PCR was performed using these primer sets from genomic DNA of all founders. Off-targeting would be Indicated by a new band at 850bp, indicating insertion of the tags in the off-target locus. No off-targeting was detected by this method (below).

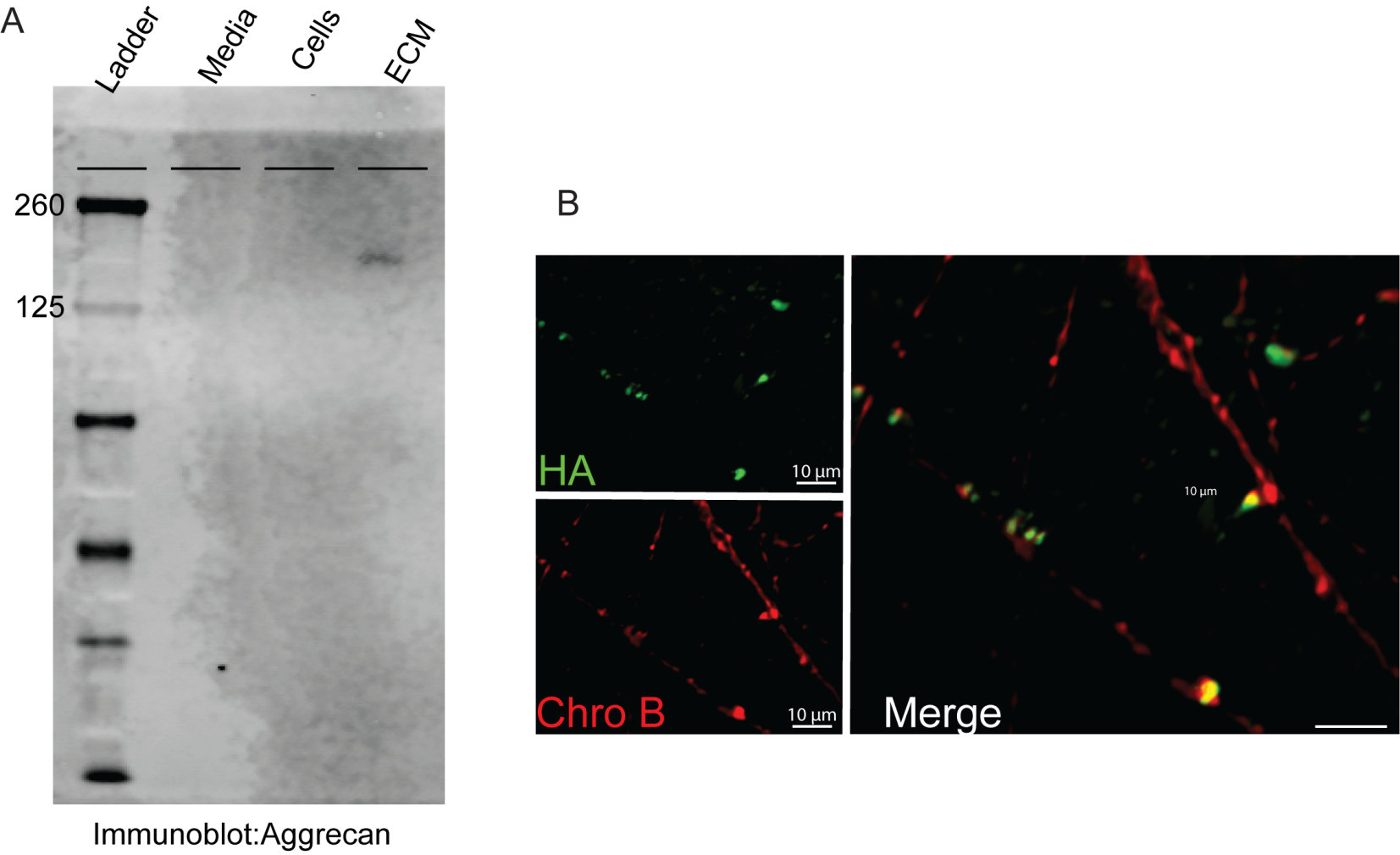

Fig. S2. Validation of intracellular/extracellular (media)/extracellular(ECM) fractionation protocol

**A**) The ECM marker protein Aggrecan is found in the ECM fraction, but absent from cell lysate and media protein fractions. **B**) Immunocytochemistry of *Pants^SOG-Y^* primary neurons co-stained with antibodies to HA and the LDCV marker chromogranin B.

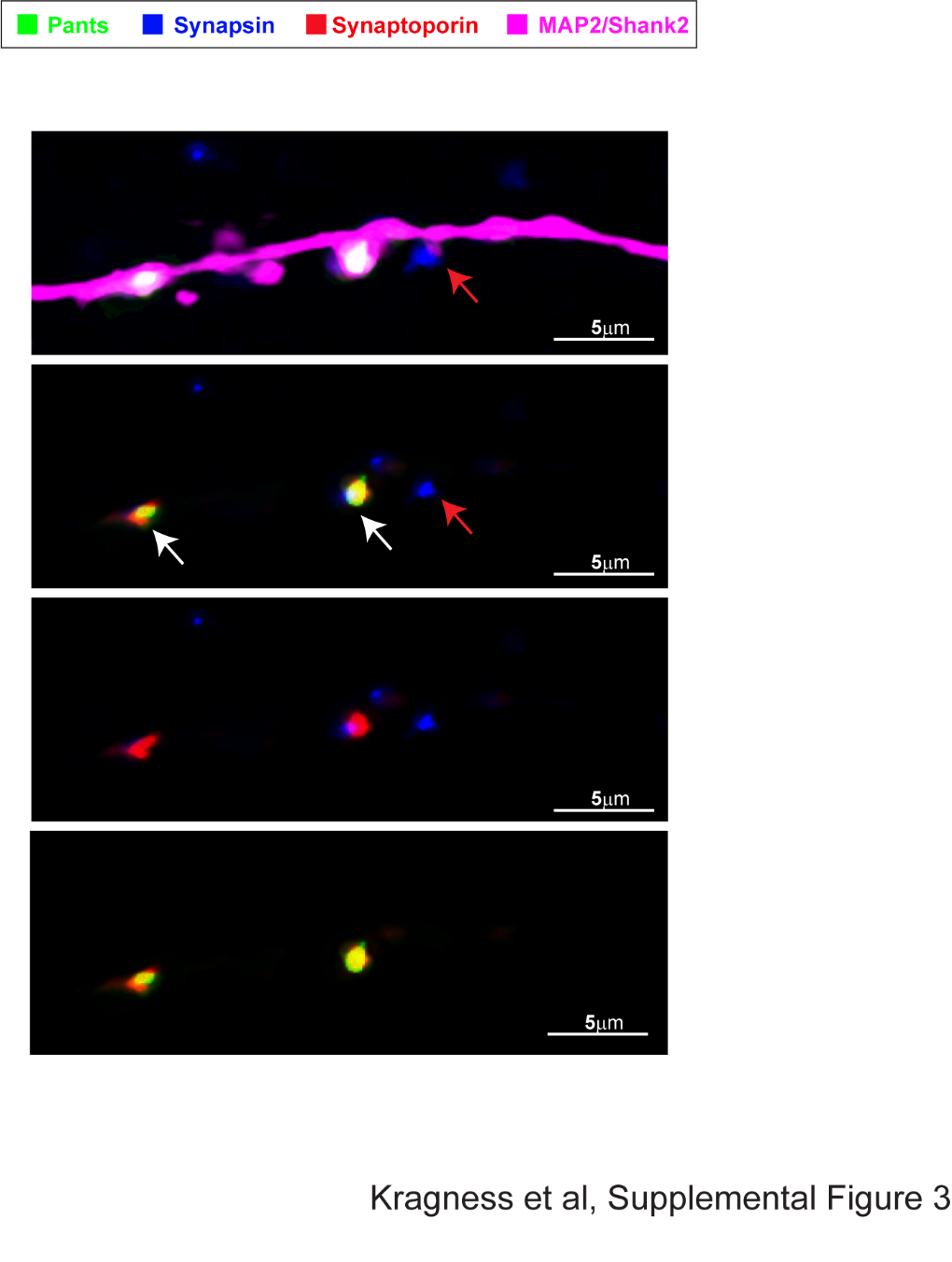

**Fig. S3: Complex spines observed on cultured hippocampal neurons are positive for synaptoporin, a marker for thorny excrescences**. Two morphologically complex spines (white arrows) are positive for Pants (green) and for the TE marker synaptoporin (red), while a mushroom spine (red arrow) is not.

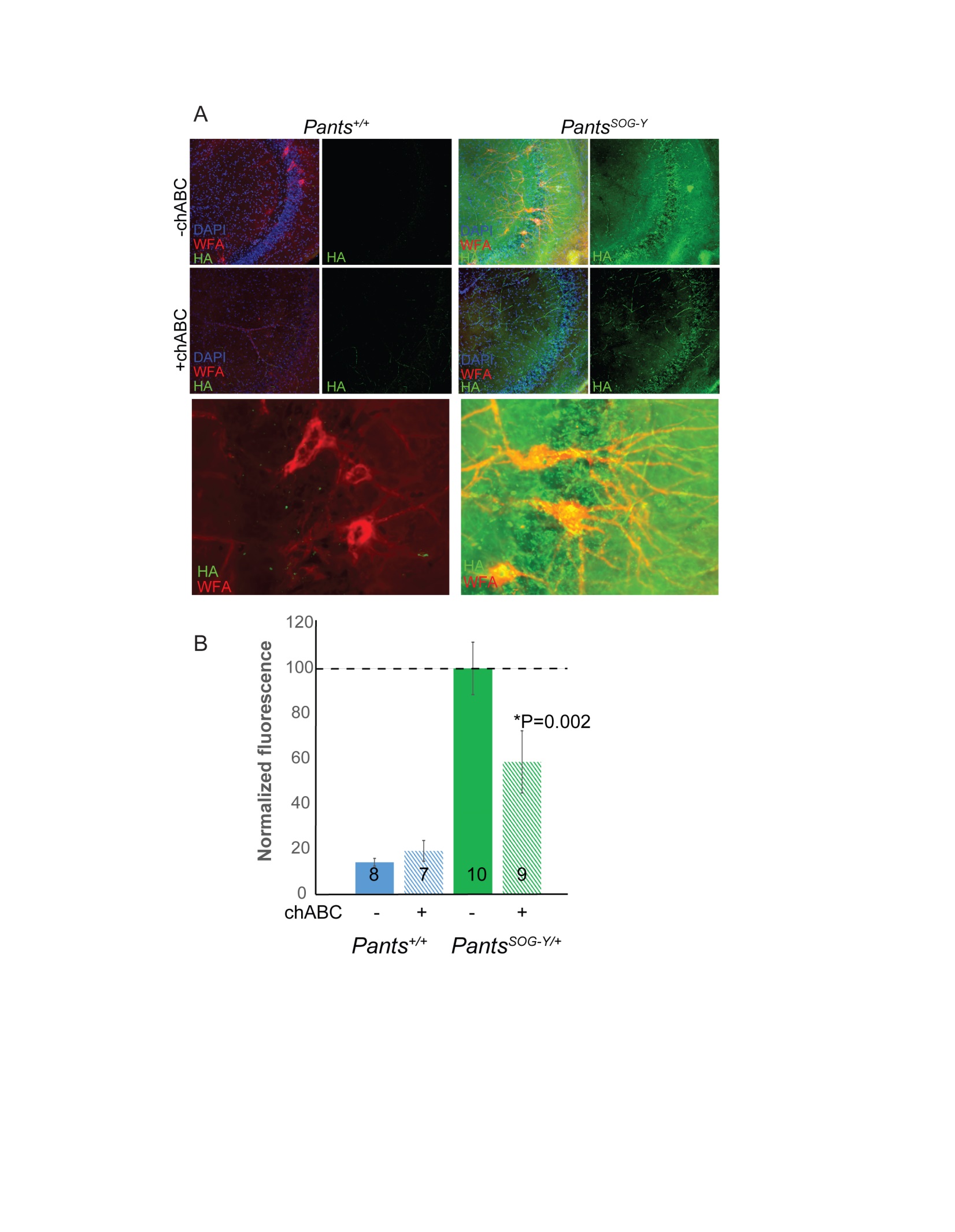

**Fig. S4. Pants is extra-synaptic in the adult hippocampus.** **A)** Immunohistochemical co-localization of HA with WFA, an ECM marker in 16 week hippocampal area CA3 in the absence and presence of chondroitinase ABC, and enzyme that digests ECM. **B)** Quantification of fluorescence intensity of HA staining in area CA3 in the absence and presence of chABC (n = 3 mice each condition; number of slices on each bar).

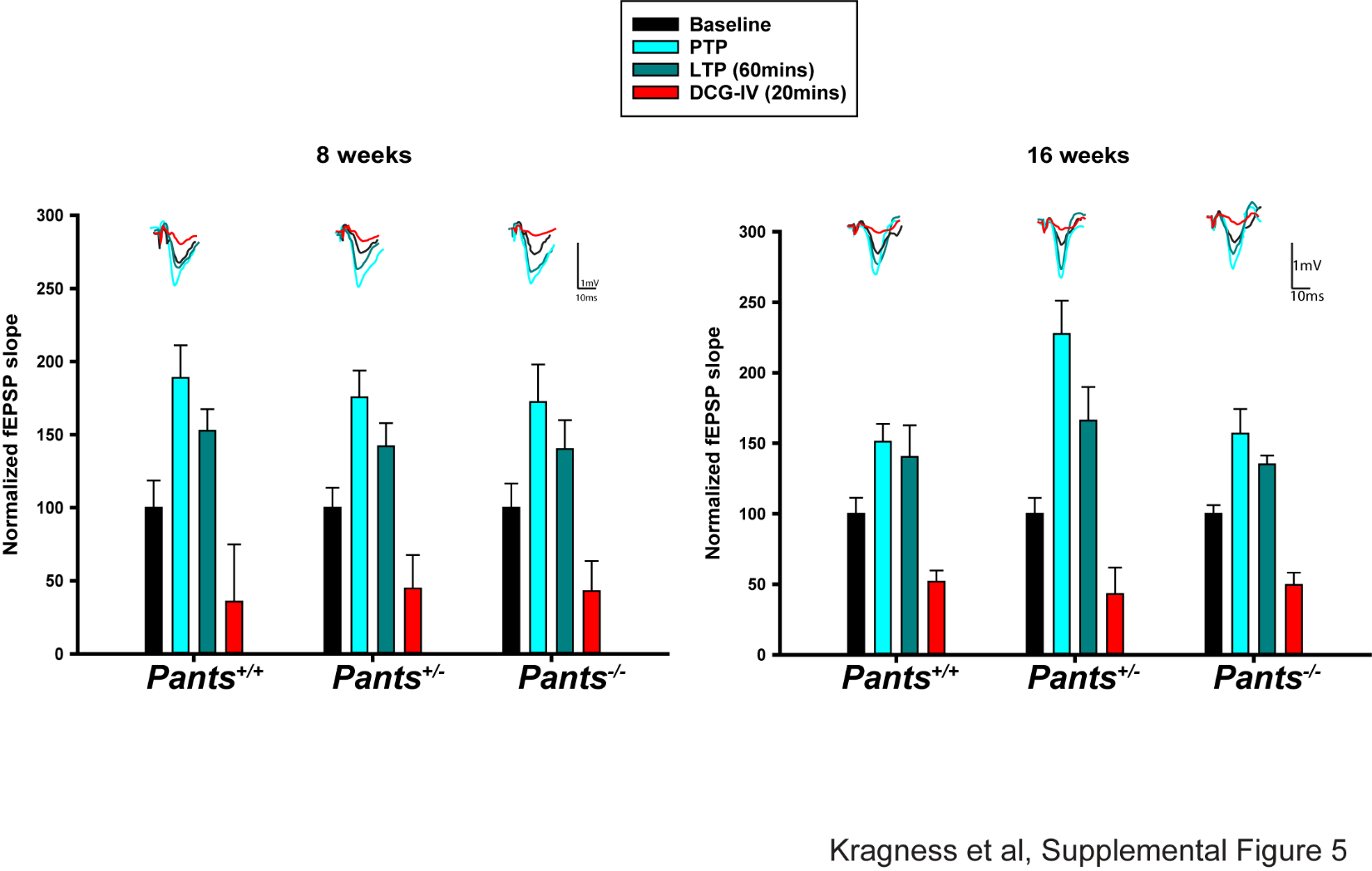

**Fig. S5. Verification of Mossy Fiber recordings by DCG-IV wash-in.** Average slope of field responses at baseline and post-tetanic potentiation and LTP for all 3 genotypes. Red bars indicate field responses after DCG-IV wash-in. Abolishment of LTP by this blocker is a specific feature of MF LTP.

| **Table S1: Antibodies/Stains Used** | | | | |
| --- | --- | --- | --- | --- |
| **Antigen** | **Host** | **Manufacturer** | **Catolog#** | **Uses and Dilutions** |
| Map2 | Chicken | Invitrogen | PA1-10005 | ICC 1:1,000 |
| Shank2 | Guinea Pig | SySy | 162 204 | ICC 1:300 |
| Synapsin 1a/b | Goat | Santa Cruz | sc-7379 | ICC 1:500 |
| β-Actin | Mouse | Sigma-Aldrich | A5316 | WB 1:10,000 |
| HA | Mouse | Santa Cruz | sc-7392 | ICC 1:300 |
| HA | Rabbit | Cell Signaling Technologies | 3724S | WB 1:500 ICC 1:300 |
| Rtn4/Nogo-A | Rabbit | Thermo Fisher Sci | PA520366 | WB 1:1,000 ICC 1:50 |
| Ncam1 | Rabbit | Sigma-Aldrich | AB5032 | WB 1:1,000 |
| ChgB | Rabbit | SySy | 259 103 | ICC 1:200 |
| Ranbp2 | Mouse | Santa Cruz | sc-74518 | WB 1:500 |
| CCT8 | Rabbit | Proteintech | 12263-1-AP | WB 1: 500 |
| Streptavidin-HRP | n/a | Invitrogen | S911 | WB 0.3μg/ml |
| WFA | n/a | Vector Labs | B-1355-2 | ICC 1:250 |
| GluR1 | Mouse | Thermo Fisher Sci | MA527694 | ICC 1:100 |
| Pants | Rabbit | Pierce | N/A | ICC 1:50 |
| Synaptoporin | Rabbit | SySy | 102 002 | ICC 1:200 |
| V5 | Rabbit | Cell Signaling Technologies | D38HQ | WB 1:1000 |

Table S1.

Information about antibodies used in the study.

Table S2. Unique proteins biotinylated in Pants-Turbo transgenic cell line

| **Accession** | **Description** | **Score Sequest HT** | **Gene Symbol** | **Involved in Plasticity/ memory** | **Implicated in Cognitive Disease** |
| --- | --- | --- | --- | --- | --- |
| Q9ERU9 | E3 SUMO-protein ligase RanBP2 [OS=Mus musculus] | 26.94 | Ranbp2 |  |  |
| P13595 | Neural cell adhesion molecule 1 [OS=Mus musculus] | 26.29 | Ncam1 | YES (*9-12*) |  |
| P46660 | alpha-internexin [OS=Mus musculus] | 10.97 | Ina |  | YES |
| Q62448 | Eukaryotic translation initiation factor 4 gamma 2 [OS=Mus musculus] | 9.02 | Eif4g2 |  |  |
| P42932 | T-complex protein 1 subunit theta [OS=Mus musculus] | 8.58 | Cct8 |  |  |
| Q8BGD9 | eukaryotic translation initiation factor 4B [OS=Mus musculus] | 8.32 | Eif4b |  | YES (*13*) |
| Q9D9T8 | EF-hand domain-containing protein 1 [OS=Mus musculus] | 6.63 | Efhc1 |  |  |
| Q3UHK8 | Trinucleotide repeat-containing gene 6A protein [OS=Mus musculus] | 5.72 | Tnrc6a |  |  |
| Q99P72-2 | Reticulon-4 [OS=Mus musculus] | 5.29 | Rtn4 | YES (*14, 15*) | YES (*16-21*) |
| Q9QZE5 | Coatomer subunit gamma-1 [OS=Mus musculus] | 4.79 | Copg; Copg1 |  |  |
| Q8BTI8 | serine/arginine repetitive matrix protein 2 [OS=Mus musculus] | 4.6 | Srrm2 |  | YES (*22, 23*) |
| G3X9C2 | F-box only protein 50 [OS=Mus musculus] | 4.42 | Nccrp1 |  |  |
| P70302 | stromal interaction molecule 1 [OS=Mus musculus] | 4.3 | Stim1 | YES (*24-26*) |  |
| Q80ZM8 | Cardiolipin synthase (CMP-forming) [OS=Mus musculus] | 4.04 | Crls1 |  |  |
| Q8BHA3-2 | Isoform 2 of Putative D-tyrosyl-tRNA(Tyr) deacylase 2 [OS=Mus musculus] | 3.49 | 6530401N04Rik; Dtd2 |  |  |
| Q3TLH4 | Protein Prrc2c [OS=Mus musculus] | 3.34 | Prrc2c |  |  |
| Q7TPV4 | Myb-binding protein 1A [OS=Mus musculus] | 2.95 | Mybbp1a |  |  |
| P55821 | Stathmin-2 [OS=Mus musculus] | 2.61 | Stmn2 | YES (*27-29*) | YES (*30, 31*) |
| G3X939 | sodium/hydrogen exchanger 3 [OS=Mus musculus] | 2.36 | Slc9a3 |  |  |
| Q3UKK2 | Carcinoembryonic antigen-related cell adhesion molecule 5 [OS=Mus musculus] | 2.34 | Ceacam5 |  |  |
| Q9WTQ5-1 | A-kinase anchor protein 12 [OS=Mus musculus] | 2.32 | Akap12 | YES (*32, 33*) |  |
| Q5BLK4 | terminal uridylyltransferase 7 [OS=Mus musculus] | 2.32 | Zcchc6 |  |  |
| O35129 | Prohibitin-2 [OS=Mus musculus] | 2.16 | Phb2 |  |  |
| Q99KX1 | Myeloid leukemia factor 2 [OS=Mus musculus] | 2.08 | Mlf2 |  |  |
| Q6A065 | Centrosomal protein of 170 kDa [OS=Mus musculus] | 1.95 | Cep170 |  |  |
| P24270 | catalase [OS=Mus musculus] | 1.94 | Cat |  |  |
| Q80TB8 | Synaptic vesicle membrane protein VAT-1 homolog-like [OS=Mus musculus] | 1.92 | Vat1l |  |  |
| Q05816 | Fatty acid-binding protein, epidermal [OS=Mus musculus] | 1.9 | Fabp5 |  | YES (*34, 35*) |

Data Supp_Table_S3

Raw proteins identified in streptavidin pulldown from N2a negative control

Data Supp_Table_S4

Raw proteins identified in streptavidin pulldown from Pants-TurboID transgenic cell line. Contaminant value of False indicates that the protein is unique to the transgenic samples.
